# Intragenic Recombination Influences Rotavirus Diversity and Evolution

**DOI:** 10.1101/794826

**Authors:** Irene Hoxie, John J. Dennehy

## Abstract

Because of their replication mode and segmented dsRNA genome, homologous recombination is assumed to be rare in the rotaviruses. We analyzed 23,627 complete rotavirus genome sequences available in the NCBI Virus Variation database, and found 109 instances of homologous recombination, at least 11 of which prevailed across multiple sequenced isolates. In one case, recombination may have generated a novel rotavirus VP1 lineage. We also found strong evidence for intergenotypic recombination in which more than one sequence strongly supported the same event, particularly between different genotypes of segment 9, which encodes the serotype protein, VP7. The recombined regions of many putative recombinants showed amino acid substitutions differentiating them from their major and minor parents. This finding suggests that these recombination events were not overly deleterious, since presumably these recombinants proliferated long enough to acquire adaptive mutations in their recombined regions. Protein structural predictions indicated that, despite the sometimes substantial amino acid replacements resulting from recombination, the overall protein structures remained relatively unaffected. Notably, recombination junctions appear to occur non-randomly with hot spots corresponding to secondary RNA structures, a pattern seen consistently across segments. In total, we found strong evidence for recombination in nine of eleven rotavirus A segments. Only segment 7 (NSP3) and segment 11 (NSP5) did not show strong evidence of recombination. Collectively, the results of our computational analyses suggest that, contrary to the prevailing sentiment, recombination may be a significant driver of rotavirus evolution and may influence circulating strain diversity.

## Introduction

The non-enveloped, dsRNA rotaviruses of the family *Reoviridae* are a common cause of acute gastroenteritis in young individuals of many bird and mammal species (Desselberger 2014). The rotavirus genome consists of 11 segments, each coding for a single protein with the exception of segment 11, which encodes two proteins, NSP5 and NSP6 (Desselberger 2014). Six of the proteins are structural proteins (VP1-4, VP6 and VP7), and the remainder are non-structural proteins (NSP1-6). The infectious virion is a triple-layered particle consisting of two outer-layer proteins, VP4 and VP7, a middle layer protein, VP6, and an inner capsid protein, VP2. The RNA polymerase (VP1) and the capping-enzyme (VP3) are attached to the inner capsid protein. For the virus to be infectious (at least when not infecting as an extracellular vesicle), the VP4 spike protein must be cleaved by a protease, which results in the proteins VP5* and VP8* (Arias, Romero et al. 1996). Because they comprise the outer layer of the virion, VP7 and VP4 are capable of eliciting neutralizing antibodies, and are used to define G (glycoprotein) and P (protease sensitive) serotypes respectively (Matthijnssens, Ciarlet et al. 2008, Nair, Feng et al. 2017). Consequently, VP7 and VP4 are likely to be under strong selection for diversification to mediate cell entry or escape host immune responses (McDonald, Matthijnssens et al. 2009, Kirkwood 2010, Patton 2012).

Based on sequence identity and antigenic properties of VP6, 10 different rotavirus groups (A–J) have been identified, with group A rotaviruses being the most common cause of human infections (Matthijnssens, Otto et al. 2012, Mihalov-Kovacs, Gellert et al. 2015, Banyai, Kemenesi et al. 2017). A genome classification system based on established nucleotide percent cutoff values has been developed for group A rotaviruses (Matthijnssens, Ciarlet et al. 2008, Matthijnssens, Ciarlet et al. 2011). In the classification system, the segments VP7-VP4-VP6-VP1-VP2-VP3-NSP1-NSP2-NSP3-NSP4-NSP5/6 are represented by the indicators Gx-P[x]-Ix-Rx-Cx-Mx-Ax-Nx-Tx-Ex-Hx, (x = Arabic numbers starting from 1), respectively (Matthijnssens, Ciarlet et al. 2008, Matthijnssens, Ciarlet et al. 2011). To date, between 20 to 51 different genotypes have been identified for each segment, including 51 different VP4 genotypes (P[1]–P[51]) and 36 different VP7 genotypes (G1–G36), both at 80% nucleotide identity cutoff values (Steger, Boudreaux et al. 2019).

The propensity of rotavirus for coinfection and outcrossing with other rotavirus strains makes it a difficult pathogen to control and surveil, even with current vaccines (Rahman, Matthijnssens et al. 2007, Matthijnssens, Ciarlet et al. 2008, Matthijnssens, Bilcke et al. 2009, Kirkwood 2010, Ghosh and Kobayashi 2011, Sadiq, Bostan et al. 2018). Understanding rotaviral diversity expansion, genetic exchange between strains (especially between the clinically significant type I and type II genogroups), and evolutionary dynamics resulting from coinfections have important implications for disease control (Rahman, Matthijnssens et al. 2007, Matthijnssens, Ciarlet et al. 2008, Matthijnssens, Bilcke et al. 2009, Kirkwood 2010, Ghosh and Kobayashi 2011, Sadiq, Bostan et al. 2018). Rotavirus A genomes have high mutation rates (Matthijnssens, Heylen et al. 2010, Donker and Kirkwood 2012, Sadiq, Bostan et al. 2018), undergo frequent reassortment (Ramig and Ward 1991, Ramig 1997, Ghosh and Kobayashi 2011, McDonald, Nelson et al. 2016), and the perception is that these two processes are the primary drivers of rotavirus evolution (Doro, Farkas et al. 2015, Sadiq, Bostan et al. 2018). Genome rearrangements may also contribute to rotavirus diversity, but are not believed to be a major factor in rotavirus evolution (Desselberger 1996). Homologous recombination, however, is thought to be especially rare in rotaviruses due to their dsRNA genomes (Ramig 1997, McDonald, Nelson et al. 2016, Varsani, Lefeuvre et al. 2018). Unlike +ssRNA (Lukashev 2005) viruses and DNA viruses (Pérez-Losada, Arenas et al. 2015), dsRNA viruses cannot easily undergo intragenic recombination because their genomes are not replicated in the cytoplasm by host polymerases, but rather within nucleocapsids by viral RNA-dependent RNA polymerase (Ramig 1997, Patton, Vasquez-Del Carpio et al. 2007). Genome encapsidation should significantly reduce the opportunities for template switching, the presumptive main mechanism of intramolecular recombination (Lai 1992, Pérez-Losada, Arenas et al. 2015).

Despite the expectation that recombination should be rare in rotavirus A, there are nevertheless numerous reports of recombination among rotaviruses in the literature (Suzuki, Gojobori et al. 1998, Parra, Bok et al. 2004, Phan, Okitsu et al. 2007, Phan, Okitsu et al. 2007, Cao, Barro et al. 2008, Martinez-Laso, Roman et al. 2009, Donker, Boniface et al. 2011, Jere, Mlera et al. 2011, Esona, Roy et al. 2017, Jing, Zhang et al. 2018). However, a comprehensive survey of 797 rotavirus A genomes failed to find any instances of the same recombination event in multiple samples (Woods 2015). There are two possible explanations for this outcome. Either the putative recombinants were spurious results stemming from poor analytical technique or were poorly fit recombinants that failed to increase in frequency in the population such that they would be resampled (Woods 2015). The main implications of this study are that recombination among rotavirus A is rare, usually disadvantageous, and not a significant factor in rotavirus evolution.

Since the number of publicly available rotavirus A whole segment genomes is now over 23,600, it is worth revisiting these conclusions to see if they are still valid. To this end, we used bioinformatics tools to identify possible instances of recombination among all available complete rotavirus A genome sequences available in the NCBI Virus Variation database as of May, 2019. We found strong evidence for recombination events among all rotavirus A segments with the exception of NSP3 and NSP5. In several cases, the recombinants were fixed in the population such that several hundred sampled strains showed remnants of this same event. These reports suggest that rotavirus recombination occurs more frequently than is generally appreciated, and can significantly influence rotavirus A evolution.

## Methods

### Sequence Acquisition and Metadata Curation

We downloaded all complete rotavirus genomes from NCBI’s Virus Variation Resource as of May 2019 (n = 23,627) (Hatcher, Zhdanov et al. 2017). Laboratory strains were removed from the dataset. Genomes that appeared to contain substantial insertions were excluded as well. Avian and mammalian strains were analyzed separately as recombination analyses between more divergent genomes can sometimes confound the results. No well-supported events were identified among the avian strains. For each rotavirus genome, separate fasta files were downloaded for each of the 11 segments. Metadata including host, country of isolation, collection date, and genotype (genotype cutoff values as defined by (Matthijnssens, Ciarlet et al. 2008)) were recorded. The genotype cutoff values allowed for the differentiation of events that occurred between two different genotypes— intergenotypic—from those which occurred between the same genotype—intragenotypic.

### Recombination Detection and Phylogenetic Analysis

All 11 segments of each complete genome were separately aligned using MUSCLE v3.8.31 (Edgar 2004) after removing any low quality sequence (e.g., ‘Ns’). Putative recombinants were identified using RDP4 Beta 4.97, a program, which employs multiple recombination detection methods to minimize the possibility of false positives (Martin, Murrell et al. 2015). All genomes were analyzed, with a p-value cut off < 10E-4, using 3Seq, Chimæra, SiScan, MaxChi, Bootscan, Geneconv, and RDP as implemented in RDP4 (Martin, Murrell et al. 2015). We eliminated any strains that did not show a putative recombination event predicted by at least 6 of the above listed programs. We then ran separate phylogenic analyses on the ‘major parent’ and ‘minor parent’ sequences of putative recombinants in BEAST v1.10.4 using the general time reversible (GTR) + Γ + I substitution model (Suchard, Lemey et al. 2018). ‘Parent’ in this case does not refer to the actual progenitors of the recombinant strain, but rather those members of the populations whose genome sequences most closely resemble that of the recombinant. Significant phylogenetic incongruities with high posterior probabilities between the ‘major parent’ and ‘minor parent’ sequences were interpreted as convincing evidence for recombination.

### Phylogenetic Analysis of VP1 and VP3

Segment 1 (VP1) and segment 3 (VP3) each showed evidence for recombination events resulting in a novel lineage, wherein many isolates reflected the same recombination event. To analyze these events more thoroughly, we split alignments of a subset of environmental isolates to include flagged clades, minor parent clades, and major parent clades along with outgroup clades to improve accuracy of tip dating. We generated minor and major parent phylogenies using BEAST v1.10.4 (Suchard, Lemey et al. 2018). We used tip dating to calibrate molecular clocks and generate time-scaled phylogenies. The analyses were run under an uncorrelated relaxed clock model using a time-aware Gaussian Markov random field Bayesian skyride tree prior (Minin, Bloomquist et al. 2008). The alignments were run using a GTR + Γ + I substitution model and partitioned by codon position. Log files in Tracer v1.7.1 (Rambaut, Drummond et al. 2018) were analyzed to confirm sufficient effective sample size (ESS) values, and trees were annotated using a 20% burn in. The alignments were run for three chains with a 200,000,000 Markov chain Monte Carlo (MCMC) chain length, analyzed on Tracer v1.7.1 (Rambaut, Drummond et al. 2018), and combined using LogCombiner v1.10.4 (Drummond and Rambaut 2007). Trees were annotated with a 10% burn, and run in TreeAnnotator v1.10.4 (Suchard, Lemey et al. 2018). The best tree was visualized using FigTree v1.4.4 (Rambaut 2019) with the nodes labeled with posterior probabilities.

### RNA Secondary Structure Analysis

To test the hypothesis that recombination junctions were associated with RNA secondary structure, we generated consensus secondary RNA structures of different segments and genotypes using RNAalifold in the ViennaRNA package, version 2.4.11 (Bernhart, Hofacker et al. 2008, Lorenz, Bernhart et al. 2011). The consensus structures were visualized and mountain plots were generated to identify conserved structures that may correspond to breakpoint locations. We made separate consensus alignments in two ways, by using the same genotype and by combining genotypes. Due to the high sequence variability of the observed recombinants, particularly in segments 4 (VP4) and 9 (VP7), we note that the consensus structures may vary substantially depending on the sequences used in the alignments. Segments 7 (NSP3), 10 (NSP4), and 11 (NSP5) were not analyzed due to only having one or no recombination events. Consensus structures could not reliably be made for segment 2 (VP2). While the VP2 protein is relatively conserved across genotypes, it contains insertions particularly in the C1 genotype, yet shows recombination across C1 and C2 genotypes.

### Protein Structure and Antigenic Epitope Predictions

To evaluate whether recombination events resulted in substantial (deleterious) protein structure changes, we employed LOMETS2 (Local Meta-Threading-Server) I-TASSER (Iterative Threading Assembly Refinement) (Zhang 2008, Roy, Kucukural et al. 2010, Yang and Zhang 2015) to predict secondary and tertiary protein structures. I-TASSER generates a confidence (C) score for estimating the quality of the protein models. To determine if any of the putative recombinants possessed recombined regions containing epitopes, we analyzed the amino acid sequences of all VP4, VP6, and VP7 recombinants and their major and minor parents. We used the Immune Epitope Database (IEDB) (Vita, Mahajan et al. 2019) and SVMTriP (Yao, Zhang et al. 2012) to predict conserved epitopes.

## Results

### Strong Evidence for Homologous Recombination in Rotavirus A

We identified 109 putative recombination events (identified by 6/7 RDP4 programs; Table 1, Supplementary Table 1). Of these, 67 recombination events were strongly supported, meaning they were detected by 7/7 RDP4 programs with a p-value cut off < 10E-9 (Table 1). Most recombination events detected were observed in sequences uploaded to the Virus Variation database since the last large scale analysis (Woods 2015) so differences between these results and prior studies may simply reflect the recent increase in available genome sequences.

**Table 1.** Recombination events identified among all mammalian rotavirus A genome sequences downloaded from NCBI Virus Variation database in May 2019 (Hatcher, Zhdanov et al. 2017). _a_6/7 programs implemented in RDP4 identified putative recombination event (see Methods). _b_Events identified by 7/7 programs implemented in RDP4 or events where more than one environmental isolate showed the same event (see Methods). _c_Strongly supported events (Row 3) divided by number of sequences analyzed (Row 1). _d_Number of intergenotypic events out of all putative recombination events (Row 2).

|  | VP1 | VP2 | VP3 | VP4 | VP6 | VP7 | NSP1 | NSP2 | NSP3 | NSP4 | NSP5 |
| --- | --- | --- | --- | --- | --- | --- | --- | --- | --- | --- | --- |
| **Number of sequences analyzed** | 1710 | 1600 | 1905 | 1990 | 2176 | 3887 | 1962 | 2186 | 1881 | 2430 | 1900 |
| **Putative recombination events<sup>a</sup>** | 15 | 13 | 16 | 11 | 11 | 24 | 11 | 4 | 0 | 3 | 1 |
| **Strongly supported events<sup>b</sup>** | 7 | 8 | 15 | 7 | 6 | 14 | 7 | 1 | 0 | 1 | 1 |
| **Recombination frequency<sup>c</sup>** | 4.1E-3 | 5.0E-3 | 7.9E-3 | 3.5E-3 | 2.8E-3 | 3.6E-3 | 3.6E-3 | 4.6E-4 | -- | 4.1E-4 | 5.3E-4 |
| **Intergenotypic recombination ratio<sup>d</sup>** | 2/15 | 8/13 | 4/16 | 6/11 | 7/11 | 22/24 | 3/11 | 2/4 | -- | 2/3 | 1/1 |

### Putative Recombination Events Observed in Multiple Environmental Isolates

Eleven of the recombination events identified in Table 1 were observed in more than one environmental isolate (Fig. 1; Table 2). The observation of multiple sequenced strains with the same recombination event is strong evidence that the observed event was not spurious, and was not a consequence of improper analytical technique or experimental error. Assuming the events are not spurious, there are only two ways that multiple sequenced isolates will show the same recombinant genotype. Either multiple recombination events with the same exact breakpoints occurred at approximately the same time, or the event happened once and descendants of the recombined genotype were subsequently isolated from additional infected hosts. The latter scenario is more parsimonious, and suggests that the new genotype was not reproductively impaired. Indeed, some recombined genotypes may be more fit than their predecessors, but this outcome would need to be experimentally validated with infectivity assays.

**Figure 1.**
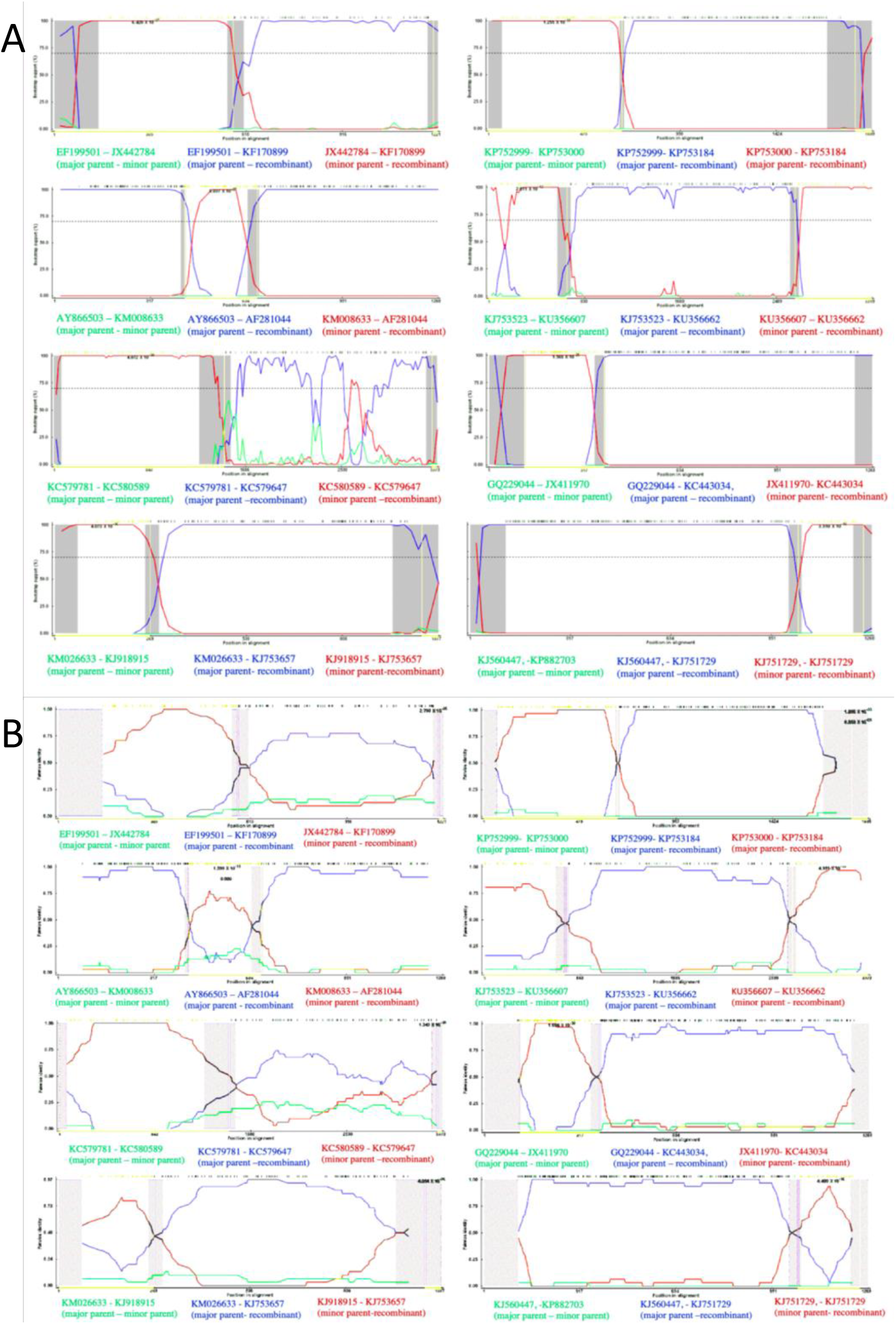
Bootscan (A) and RDP (B) analyses of putative recombinants where multiple environmental samples supported the event. From top left to bottom left: a G6-G6 event in VP7, a G9-G1 event, an R1-R1 event in VP1, and an N1-N1 event in NSP2. From top right to bottom right: an A8-A8 event in NSP1, an R2-R2 event in VP1, a G2-G1 event in VP7, and a G3-G1 event.

**Table 2.** Recombination events observed in multiple independent environmental isolates (i.e., isolates from different patients and/or sequenced by different laboratories). _a_99% confidence intervals. _b_See Fig. 2.

| Recombined Segment | Genotype(s) Involved | Number of Independent Isolates | Corrected and Uncorrected Multiple Comparisons | Breakpoints<sup>a</sup> | Accession Numbers |
| --- | --- | --- | --- | --- | --- |
| Segment 1 (VP1) | R1 | 2 | 4.787E-18 and 1.349E-10 | 35 – 1448 (1290 – 1498) | KC579647<br>KC580176 |
| Segment 1 (VP1) | R2 | Flagged in 285 samples. | 1.305E-16 and 3.332E-09 | (541-711) – (1327-1445) | KU199270<sup>b</sup> |
| Segment 1 (VP1) | R2 | 2 | 1.756E-20 and 4.951E-13 | 656 (541-711) – 2661 (2600-2735) | KU356662<br>KU356640 |
| Segment 3 (VP3) | M2 | Flagged in 551 samples. | 1.78E-12 and 4.982E-05 | 1258 (872-1389) – 1798 (1662-1917) | KX655453 |
| Segment 3 (VP3) | M1 | Flagged in 107 samples. | 2.491E-20 and 6.946E-13 | 2158 (2129-2174) – 2531 (undetermined) | KJ919553<br>KJ919517<br>KJ919551<br>KJ753665<br>JQ069727 |
| Segment 7 (NSP1) | A8 (porcine) | 3 | 3.01E-37 and 8.622E-30 | (1434-25) – (573-596) | KP753174<br>KJ753184<br>KP752951 |
| Segment 8 (NSP2) | N1 | 3 | 3.689E-13 and 4.054E-06 | 485 (455-499) – 889 (868-914) | KJ753657<br>KM026663<br>KM026664 |
| Segment 9 (VP7) | G6 (bovine) | 2 | 6.240E-12 and 2.780E-05 | 1048 (1032-128) – 481 (448-506) | HM591496<br>KF170899 |
| Segment 9 (VP7) | G1 & G2 | 2 | 5.507E-38 and 1.664E-30 | 55 (994-60) – 291 (262-296) | KC443034<br>MG181727 |
| Segment 9 (VP7) | G1 & G3 | 2 | 1.4E-23 and 4.230E-16 | 857 (833-859) – 1019 (991-51) | KJ751729<br>KP752817 |
| Segment 9 (VP7) | G1 & G9 | 2 | 5.292E-19 and 1.599E-11 | 362 (346-366) – 589 (558-605) | AF281044<br>GQ433992 |

### Segment 1 Intragenotypic Recombination Resulting in a New Lineage

Segment 1 (VP1; ∼3,302 base pairs) showed evidence of a recombination event within the R2 clade that was fixed in the population, and resulted in a new lineage (highlighted clade in Fig. 2; Fig. 3; Table 2). The multiple comparison (MC) uncorrected and corrected probabilities were 1.305E-16 and 3.332E-09 respectively. The sequence most closely related to the recombined sequence was KU199270, a human isolate from Bangladesh in 2010 (Aida, Nahar et al. 2016). Phylogenetic analysis using tip calibration suggests that the recombination event occurred no later than 2000-2005 (node Cis), so if this is a true recombination event, the 2010 sequence is not the original recombinant. The recombinant region is 100% similar to an isolate also from Bangladesh in 2010 (KU248372) (Aida, Nahar et al. 2016), which also has a putative recombinant sequence in another region of its genome. The breakpoint regions (99% CI: (541-711) — (1327-1445)) may represent a potential hotspot for segment 1 recombination as multiple recombination events show breakpoints in this area (Table 2). Phylogenetic analysis of the alignment of 250 representative sequences containing only the putative recombinant region compared with the rest of the segment 1 sequence showed a consistent subclade shift within the R2 clade. The recombination events resulted in the incorporation of the following amino acid substitutions in the recombinant strains when compared to the major parent strain: 227 (K→E), 293 (D→N), 297 (K→R), 305 (N→K), and 350 (K→E).

**Figure 2.**
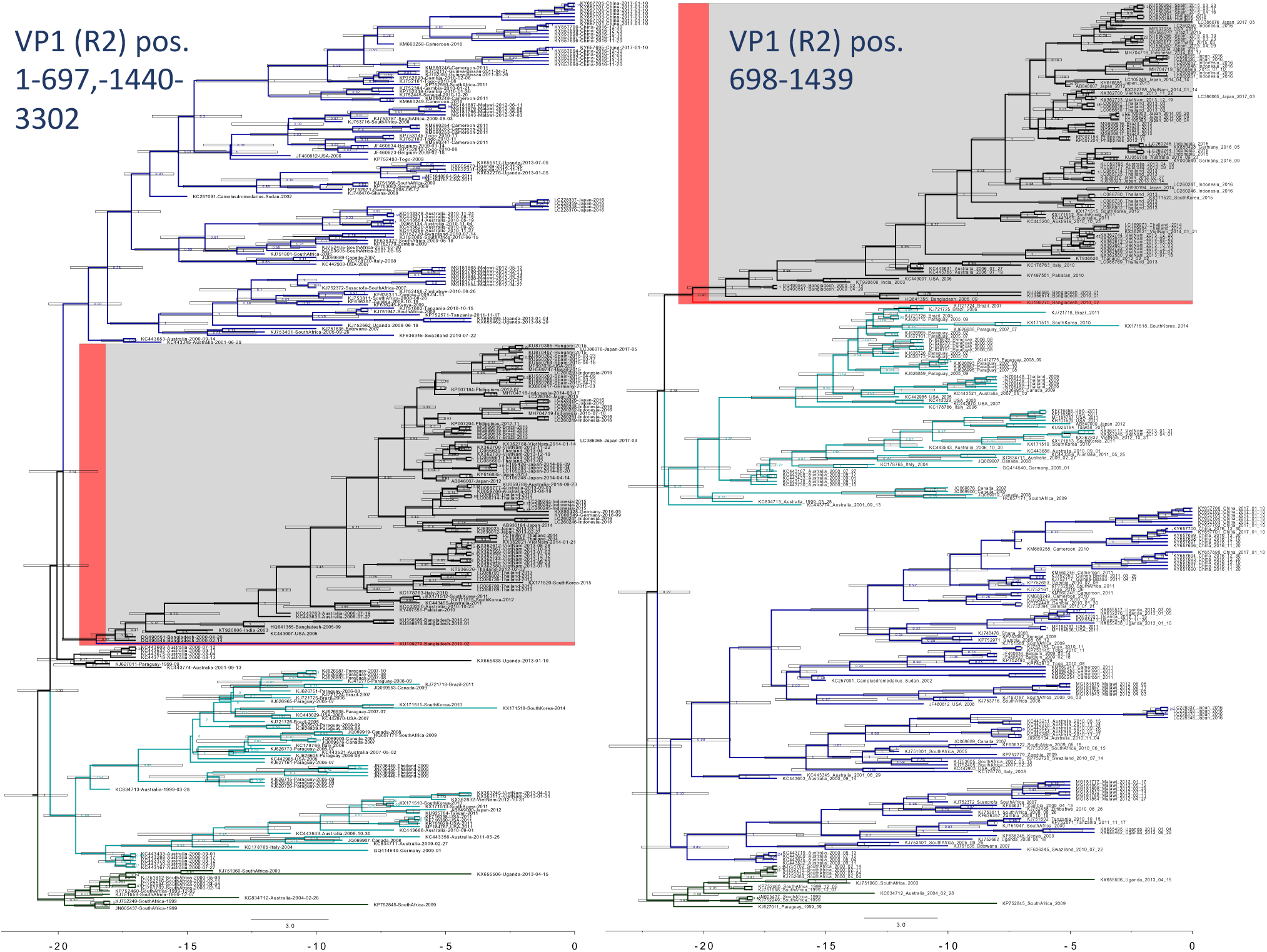
Bayesian phylogenetic analysis of a possible recombination event in VP1 that became fixed in the population. The strain closest to the recombination event is highlighted in red. Major clades are colored to show the phylogenetic incongruity. Nodes are labeled with posterior probabilities with 95% node height intervals shown. Time axis at the bottom is in years before 2017.

**Figure 3.**
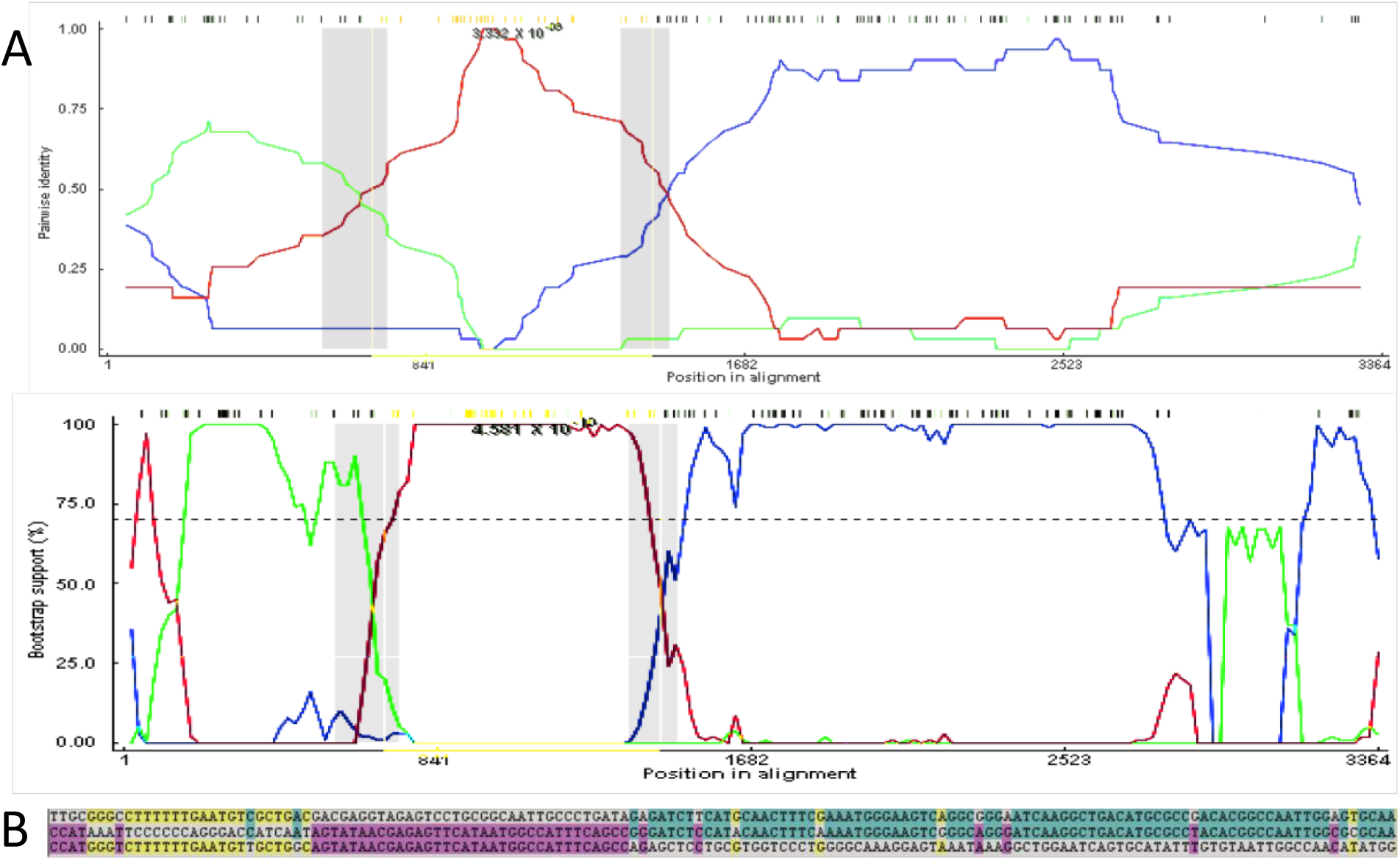
A) An RDP analysis (top) and a BootScan analysis (bottom) showing a putative recombination event between two R2 segment 1 genotypes. The red line compares the minor parent to the recombinant, blue line compares the major parent to the recombinant, and the green line compares the major parent to the minor parent. The Y-axis for RDP (top) is the pairwise identity, while the Y-axis for BootScan (bottom) is the bootstrap support. The X-axis is the sequence along segment 1. B) The relevant sites shown above color-coded to strain that the recombinant matches. The recombinant is the middle sequence, the minor parent is the bottom sequence, and the major parent is the top sequence. Mutations matching the major sequence are shown in blue, while mutations matching the minor parent are shown in purple. Yellow mutations show mutations not present in the recombinant sequence but which match the major and minor parent, possibly suggesting a second recombination event.

### Segment 3 Intragenic Recombination

Two recombination events occurring in segment 3 (VP3) appear to have fixed in the population, and are seen in many descendant sequences (Table 2). The first strain identified as a putative recombinant is KJ753665, an isolate from South Africa in 2004. The recombinant region occurs between positions (2129-2174) — 2531. This region is 98.7% identical to a porcine strain isolated in Uganda in 2016 (KY055418)(Bwogi, Jere et al. 2017), while the rest of segment 3 is 95.8% similar to a 2009 human isolate from Ethiopia (KJ752028). The Monte Carlo uncorrected and corrected probabilities were 2.491E-20 and 6.946E-13, respectively, with 107 isolates flagged as possibly derived from the recombination event. Phylogenetic analysis using 450 randomly selected VP3 sequences (excluding sequences lacking collection dates) within the putative recombinant region, along with an analysis of the genome excluding the two major recombination events, resulted in five sequences showing a significant phylogenetic incongruity (Fig. 4). The incongruity appeared between sub-lineages within the larger M1 lineage of VP3. Amino acid substitutions in the recombinant region included positions 748 (M→T) and 780 (T→M).

**Figure 4.**
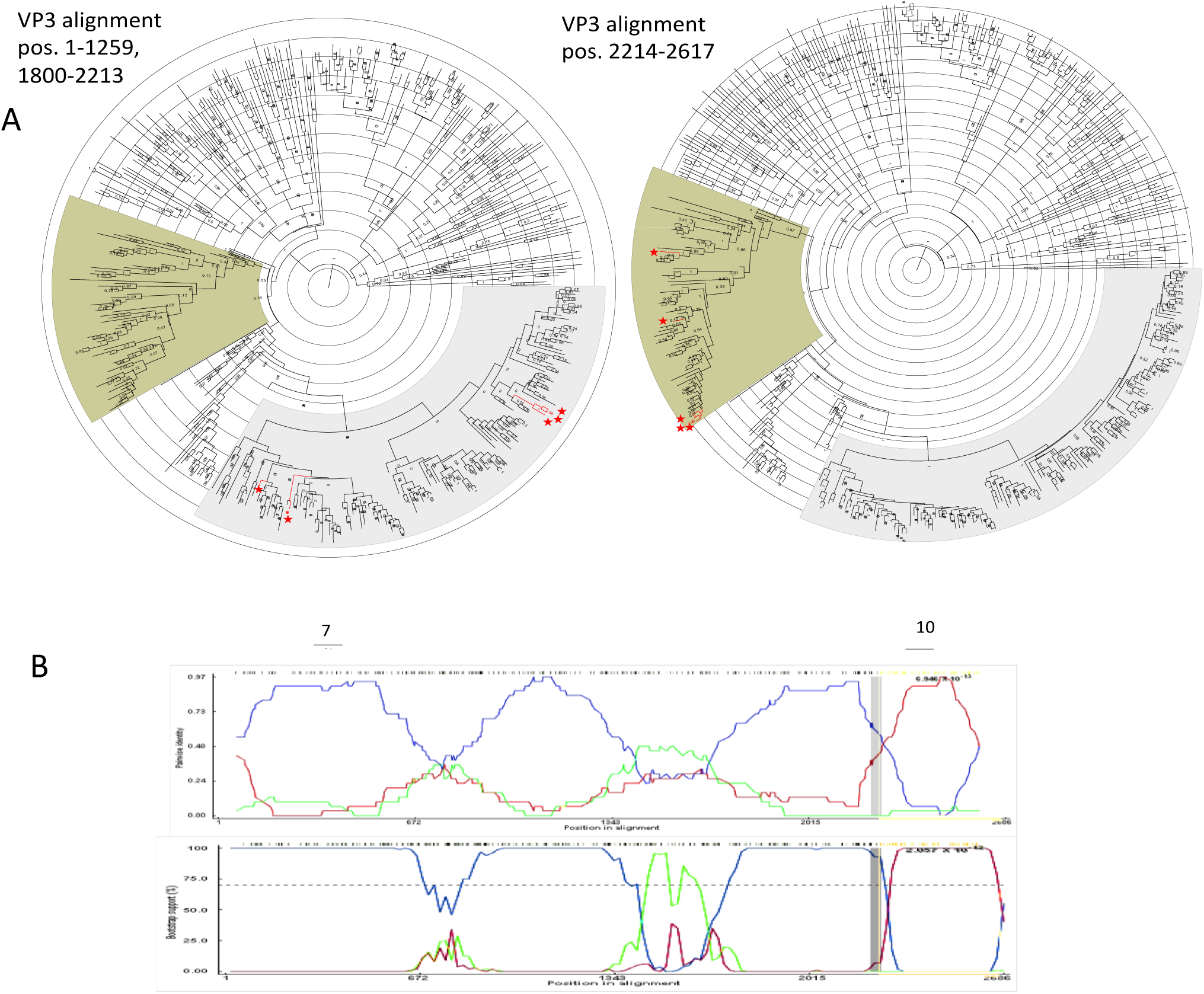
Segment 3 recombination event supported by multiple isolates. A) Phylogenetic trees made using alignments from (left) nucleotide positions 1-1259 and 1800-2213 representing the major parent region excluding second recombination event (Fig. 5), and (right) nucleotide position 2214-2617 representing the minor parent sequence. B) BootScan (top) and RDP (bottom) analyses. The red line compares the minor parent to the recombinant, blue line compares the major parent to the recombinant, and the green line compares the major parent to the minor parent. The Y-axis for the BootScan analysis (top) is the bootstrap support, while the Y-axis for the RDP analysis (bottom) is the pairwise identity. The X-axis for both analyses is the sequence along segment 3.

A second potential recombination event was identified in segment 3 between sublineages within the M2 lineage (Table 2; Fig. 5). However, when we created split alignments and ran a phylogenetic analysis in BEAST v1.10.4, only one sequence showed a phylogenetic incongruity supporting this event (KX655453). Amino acid substitutions as a result of this event included positions 405 (I→V), 412 (V→M), 414 (N→D), 441 (N→D), 458 (I→V), 459 (I→T), 468 (L→F), 473 (N→D), 486 (M→I), 518 (N→S), and 519 (E→G).

**Figure 5.**
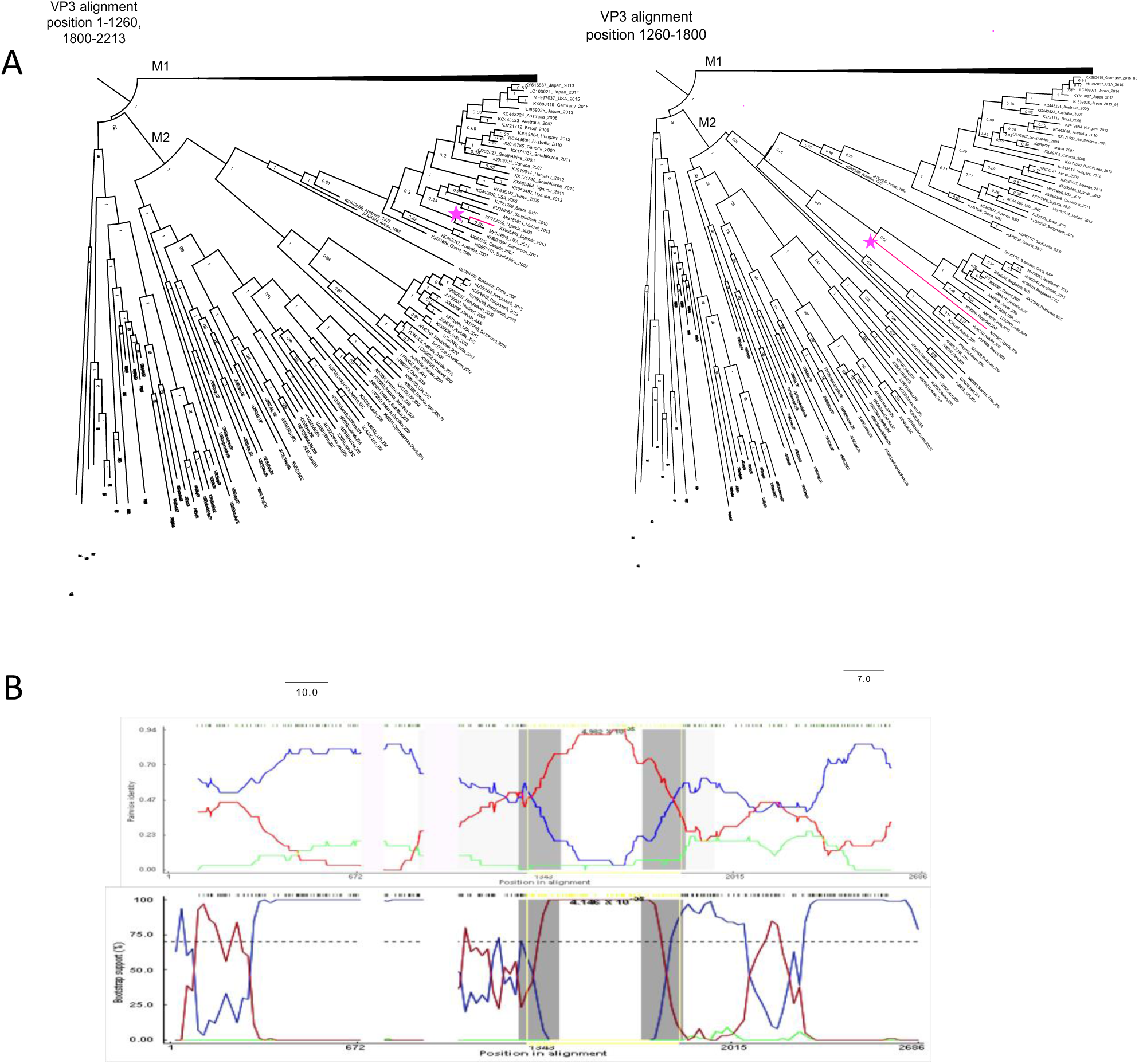
A) Phylogenetic analysis of segment 3. Left tree (major parent) was made from an alignment of nucleotide positions 1-1260 and 1900-2213 and right tree (minor parent) was made from nucleotide positions 1260-1800. M2 strains have been collapsed, and recombinant is colored in pink. B) BootScan (top) and RDP analyses (bottom of VP3 M2 putative recombinant. The red line compares the minor parent to the recombinant, blue line compares the major parent to the recombinant, and the green line compares the major parent to the minor parent. The Y-axis for BootScan is the bootstrap support, while the Y-axis for the RDP analysis is the pairwise identity. The X-axis in both analyses is the segment 3 sequence.

One possible complication with relying on phylogenetic incongruity as evidence of major recombination events is that a recombinant virus may evolve faster than the parental strains, and, despite possessing an initial fitness advantage such as immune avoidance, may eventually converge back towards the major parent’s sequence. Thus, phylogenetic analysis likely underestimates the frequency of transiently stable recombination events in a population. This phenomenon may account for the observation that the number of VP3 isolates flagged as deriving from a recombination event based on sequence analysis is greater than the prevalence of recombination predicted by the phylogenetic analysis.

### Intergenotypic Recombination

We found strong evidence for intergenotype recombination in all segments except segment 7 (NSP3) and segment 11 (NSP5) (Table 1; Supplementary Table 1). Instances of intergenotypic recombination in segment 1 (VP1) (KU714444, JQ988899) only occurred in regions where the amino acid sequence was highly conserved across genotypes, so these events resulted in few, if any, nonsynonymous mutations. Of the putative events observed in more than two environmental isolates with strong support from detection methods, only events in segment 9 (VP7) occurred between different genotypes (Table 2). The segment 9 recombinant region amino acid substitutions that match the minor parent and differ from the major parent are shown in Supplementary Table 2.

### Structural Prediction of Recombinant Proteins

The protein models generated by I-TASSER for the intergenotypic recombinant G proteins showed that, although the amino acid changes for the G1-G2 recombinant were substantial (25 changes) (Supplementary Table 2), the secondary and tertiary structures seemed largely intact (Fig. 6). The G3-G1 and G6-G6 recombinants each showed four amino acid changes. The G9-G1 recombinant showed the most secondary structure disruption, including a loss of beta sheets, a loss of antiparallel beta sheets, and slightly shorter beta sheets/helices, which could indicate lower stability. However, based on the structural modeling, the tertiary structure appeared to be maintained, suggesting that the putative recombinant glycoproteins were able to form properly folded G-proteins (Fig. 6).

**Figure 6.**
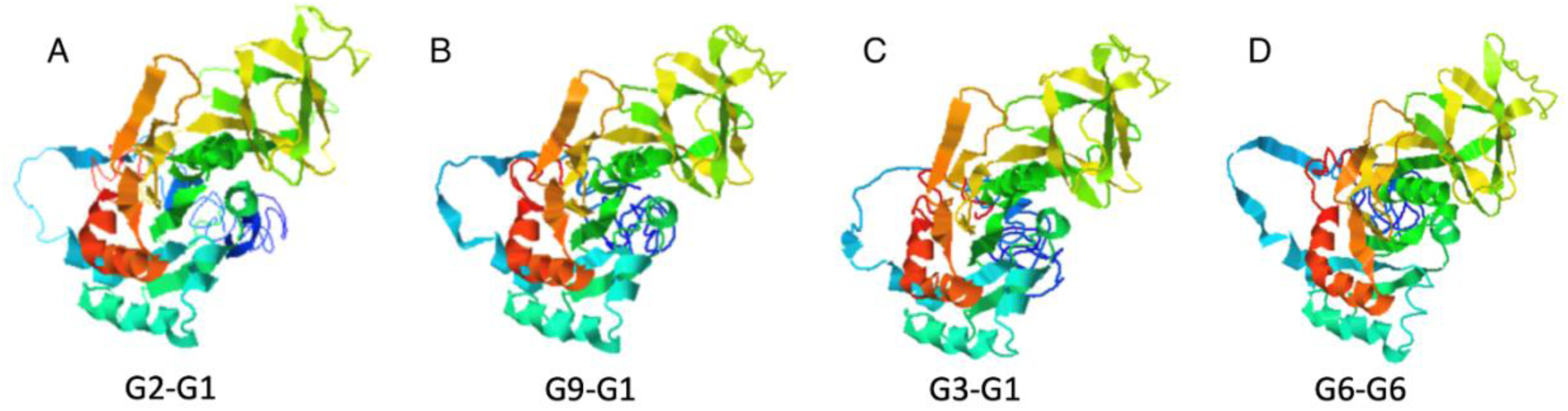
VP7 protein structures were predicted from amino acid alignments of the four strongly supported G recombinants using I-TASSER. C-scores are confidence scores estimating the quality of the predicted model, and range from [−5, 2], with higher scores indicate greater confidence A) G2-G1, C-score = −1.38, KC443034; B) G9-G1:AF281044, C-score = −1.29; C) G3-G1, C-score = −1.27 KJ751729; D) G6-G6: KF170899, C-score = −1.42.

Antigenic epitope predictions generated by IEDB (Vita, Mahajan et al. 2019) and SVMTriP (Yao, Zhang et al. 2012), as well as a large study done on mammalian G-types (Ghosh, Chattopadhyay et al. 2012), showed that VP7 recombination occasionally results in amino acid substitutions in conserved epitopes (Supplementary Table 2). For example, the amino acid sequence RVNWKKWWQV is usually flagged as, or part of, an epitope in most G types including G2, G3, G4, G6, G9, but not in G1. In the G2-G1 recombination event flagged in multiple isolates (Table 2; Figure 1), this region is altered (sometimes a KR substitution and sometimes multiple amino acid substitutions) (Supplementary Table 2). Moreover, despite containing highly variable regions, the region where the recombination occurred had low solvent accessibility. The G3-G1 recombinant had amino acid substitutions in two of four conserved epitope regions due to the recombination event. There was also a conserved epitope sequence around amino acids 297-316 in G9 proteins that was altered in the G9-G1 recombinant so it no longer appeared as an epitope.

Structural predictions generated by I-TASSER suggest that, although the amino acid sequences may diverge, the protein folding and three-dimensional structures remain relatively conserved (Fig. 6). Thus, although the amino acid sequence substitutions do not result in significant changes to the protein structure, they may nevertheless reduce binding by antibodies or T-cell receptors, and may provide a selective advantage in allowing the virus to avoid immune surveillance.

### Recombination Junctions Often Correspond to RNA Secondary Structure Elements

Breakpoint distribution plots showed the sequence regions with the most breakpoints. These breakpoints often corresponded to hairpins predicted by RNAalifold. Secondary RNA structure predictions for segment 9 (VP7) genotypes G1, G2, G6 and G9 are shown as mountain plots (Fig. 7). The breakpoints of the segment 9 recombination events correspond to areas leading to the peaks in the mountain plots (Fig. 7). The peaks indicate a conserved hairpin loop, with the sequences leading up to the peak being the double-stranded portion of the hairpin. Breakpoint distributions also appeared to correspond to secondary structure predictions made from alignments of segment 1 (VP1) and segment (VP6) (Fig. 8). Segment 4 (VP4) RNA secondary structure predictions (Supplementary Fig. 1) showed greater variation across genotypes, so while breakpoints did coincide with secondary RNA structures as predicted by the models, there were not enough events in each genotype to provide strong support that the breakpoints were correlated with secondary structure. The breakpoint distributions for NSP1 and VP3 were also non-randomly distributed across the sequence (Supplementary Fig. 1), however secondary structure predictions from these alignments were not consistent, so are not shown (Supplementary Fig. 2).

**Figure 7.**
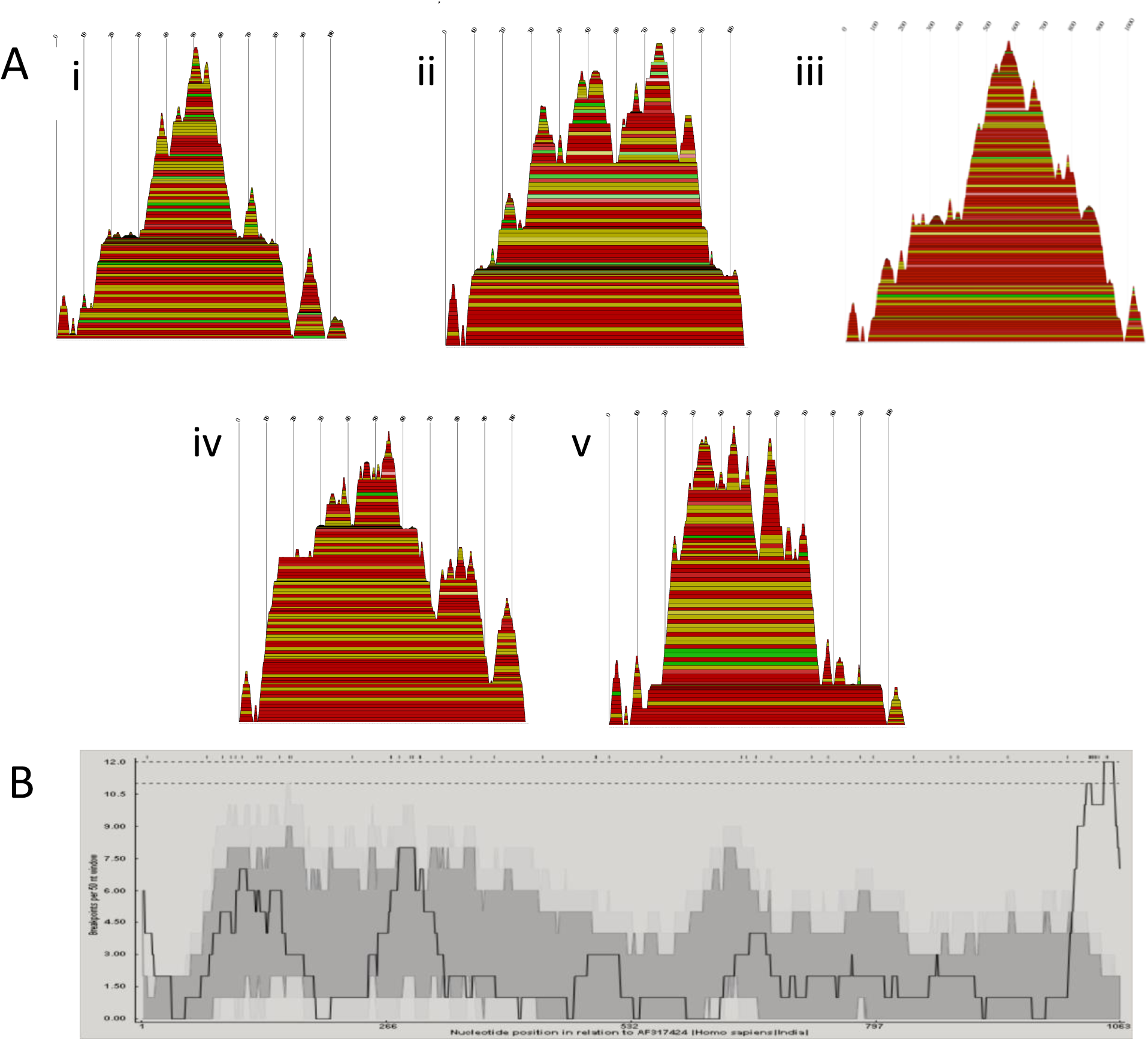
A) Consensus mountain plots made in RNAalifold using the ViennaRNA package of the predicted RNA secondary structures based on alignments made using full sequences for each of the five segment 9 G types involved in the intergenotypic recombination events: (i) G9 (ii) G6 (iii) G3 (iv) G1 (v) G2. Peaks represent hairpin loops, slopes correspond to helices, and plateaus correspond to loops. The X-axis corresponds to the sequence of the segment. Each base-pairing is represented by a horizontal box where the height of the box corresponds with the thermodynamic likelihood of the pairing. The colors correspond to the variation of base pairings at that position. Red indicates the base pairs are highly conserved across all the sequences, and black indicates the least conservation of those base pairings. B) Breakpoint distribution plots made in RDP4 of putative recombinants in segment 9. The X-axis shows the position in the sequence, and Y-axis shows the number of breakpoints per 50 nucleotide window. The highest peaks are around X = 115, 285, 1050.

**Figure 8.**
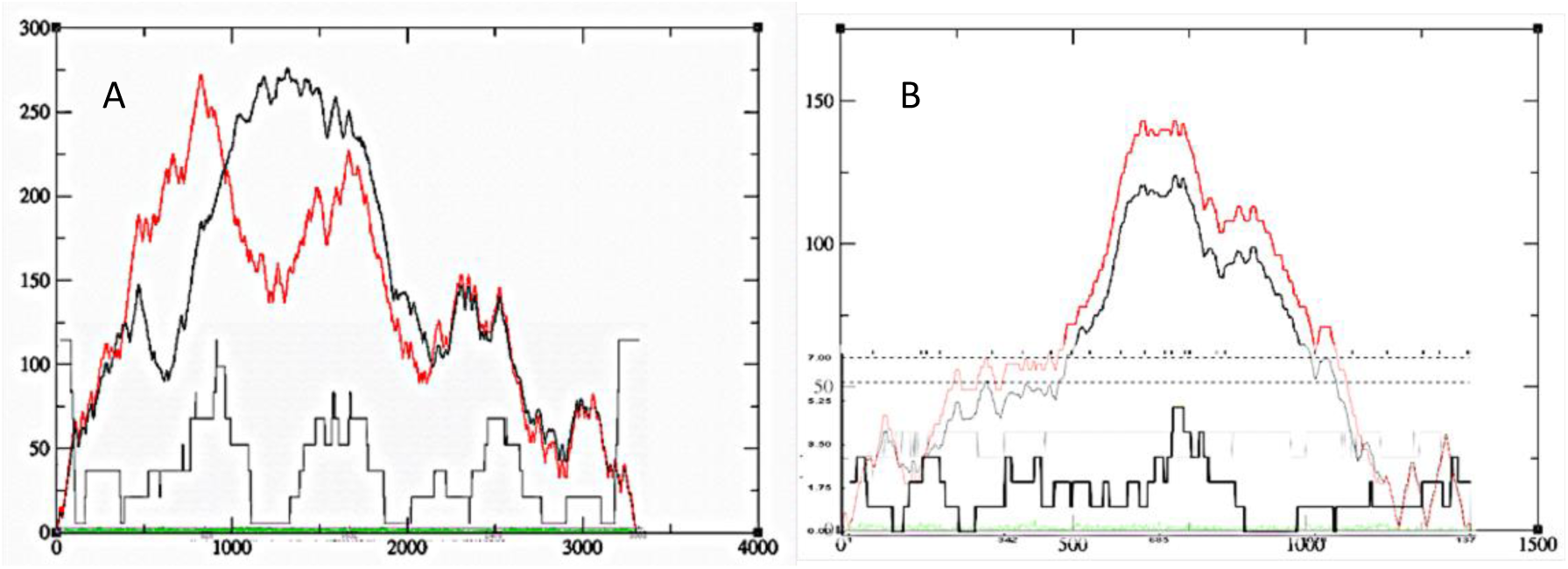
Consensus mountain plots overlaid with breakpoint distribution plots for recombination events in A) VP6 and B) VP1. Position in the nucleotide sequence is on the X-axis. The entropy curve represented in green. The black curve represents the pairing probabilities, and the red curve represents the minimum free energy structure with well-defined regions having low entropy.

## Discussion

### Detecting Recombination: Recognizing Type I and Type II Error

Apparent instances of recombination may actually be the result of convergent evolution, lineage-specific rate variation, sequencing error, poor sequence alignment, laboratory contamination or improper bioinformatics analysis (Worobey, Rambaut et al. 2002, Boni, de Jong et al. 2010, Bertrand, Töpel et al. 2012, Boni, Smith et al. 2012). Several steps can be taken to minimize incorrect attribution of viral recombination. Ideally this process should begin at the time of sequencing. First, it should be confirmed that the originating sample did not come from a host infected with multiple genotypes of the same virus type. Prior to RNA extraction, single plaques should be repeatedly picked and plated (i.e. plaque purification) to ensure that multiple genotypes are not inadvertently sequenced. Similarly, care must be taken when sequencing multiple samples of the same virus to minimize the possibility of cross-contamination.

For sequences obtained from online repositories, such precautions are rarely possible. Instead careful bioinformatics procedures can help minimize possible errors. As typical first step in identifying recombination events, virus genome sequences are analyzed with software such as RDP4 (Martin, Murrell et al. 2015), but all software programs are prone to error. For example, programs may falsely identify a recombination event when none exists (type I error) or fail to detect a true recombination event when one exists (type II error). Several studies measured errors incurred by RDP4 in the analysis of the genomes of tick-borne encephalitis virus, a positive-sense RNA flavivirus that rarely recombines (Norberg, Roth et al. 2013, Bertrand, Johansson et al. 2016). The results of the analyses indicated that recombination was overestimated in these viruses, and that certain detection methods were more prone to type I error (Norberg, Roth et al. 2013, Bertrand, Johansson et al. 2016). MaxChi, Chimaera, and SiScan showed higher false positive rates than other RDP4 programs, but had greater power to detect true recombination events. By contrast, 3Seq and GENECONV displayed lower false positive rates, but had the lowest detection power of true events.

False positives using RDP4 are especially common among closely related strains (Bertrand, Johansson et al. 2016). That said, recombination events are likely occur between closely related strains given their close spatial/temporal proximity and genetic compatibility, so caution should be used in inferring events between highly dissimilar genotypes. When a positive recombination signal has been detected, it is essential to assess its statistical significance. However, in RDP4, the *P-*value of 0.05 does not correspond to a 5% rate of false positives (Bertrand, Johansson et al. 2016), therefore we used a cut-off value of 10E-04 and focused only on events where at least six RDP4 programs detected the putative recombinant.

After identifying putative recombination events, additional strategies can be used to eliminate errors. For example, rates of type I and type II error increase with shorter length recombination regions (Boni, Zhou et al. 2008, Norberg, Roth et al. 2013, Bertrand, Johansson et al. 2016). Therefore, we ignored any putative recombination events of < 100nt with the exception of one isolate in segment 4 (NSP4) and one isolate in segment 5 (NSP5) as the putative recombinant regions in these isolates were in conserved regions at the ends of the respective segments (Supplementary Table 1). In addition, we visually inspected sequence alignments to exclude misaligned sequences (Boni, de Jong et al. 2010). Splitting alignments by major and minor parent followed by carefully parameterized BEAST runs may help distinguish genuine phylogenetic incongruity signals from spurious false positives. Furthermore, we checked for the presence of unique polymorphisms differing from the parent strains within the suspected recombination region as they may provide evidence that recombination events are not laboratory artifacts. Presumably such substitutions would reflect subsequent adaptive evolution by the recombinant virus.

In addition, we noted how many times the same recombination event occurred across multiple samples since false positive recombinants are likely to be present as single isolates in phylogenetic trees (Boni, de Jong et al. 2010). The more isolates showing the same event, the greater the probability that it represents a true recombination event, especially if the isolates were acquired and sequenced by different laboratories. Events that showed strong support, but were only isolated in one sequence are noted (Supplementary Table 1), but not discussed, as it is difficult to rule out the possibility of type I error due to PCR or mosaic contig assembly (Boni, de Jong et al. 2010, Varsani, Lefeuvre et al. 2018).

Sequence metadata can also be used to identify unlikely recombination events. For an event to be plausible, the major and minor parents should have had opportunity to coinfect the same host, which is only possible if they are congruent in time and space (Boni, de Jong et al. 2010). For example, one study identified influenza A virus strain A/Taiwan/4845/99 as a recombinant of A/Wellington/24/2000 and A/WSN/33 (He, Han et al. 2008). Given that the two parents were isolated 77 years apart in different parts of the world, it is exceedingly unlikely that is a natural recombination event. Any putative recombination events should be carefully screened to determine if the parental strains could have plausibly interacted. In Supplementary Table 1, we include information on source species, year and place of isolation, % average nucleotide identity, and genogroup for all putative recombinants and their major and minor parents.

### Naturally High Coinfection in Rotavirus A

Some features of rotavirus biology make recombination not only possible, but also relatively plausible. Rotaviruses are often released from cells as aggregates of approximately 5-15 particles contained within extracellular vesicles (Santiana, Ghosh et al. 2018). While it is not yet clear whether these extracellular vesicles can contain different rotavirus genotypes, they do allow for rotavirus coinfection even at low multiplicities of infection. Thus, the physical barriers to recombination in dsRNA viruses (Lai 1992) may be offset by the high rates of coinfection resulting from vesicle transmission of rotaviruses.

Furthermore, infection of hosts by multiple rotavirus strains appears to be relatively common. In a study of 100 children in the Detroit area, G and P typing, which identifies the serotype of the VP7 and VP4 proteins respectively, revealed that ∼10% of patients were infected with multiple rotavirus A strains (Abdel-Haq, Thomas et al. 2003). Similarly high frequencies of G and P mixed genotype infections were observed in children sampled in India (three studies showing multiple G types in 11.3 %, 12% and 21% of samples) (Husain, Seth et al. 1996, Jain, Das et al. 2001, Khetawat, Dutta et al. 2002), Spain (>11.4% of samples) (Sánchez-Fauquier, Montero et al. 2006), Kenya (5.9%) (Kiulia, Peenze et al. 2006), Africa (12%) (Mwenda, Ntoto et al. 2010), and Mexico (5.6% in 2010, 33.5% in 2012) (Anaya-Molina, De La Cruz Hernández et al. 2018). Even higher frequencies of mixed genotype infections were observed in whole genome studies. For example, among 39 Peruvian fecal samples genotyped using multiplexed PCR, 33 (84.6%) showed evidence of multiple rotavirus genotypes (Rojas, Dias et al. 2019). In another study, whole genome deep sequencing revealed that 15/61 (25%) samples obtained in Kenya contained multiple rotavirus genotypes (Mwanga, Nyaigoti et al. 2018). Given the high genetic diversity of rotavirus populations (Kirkwood 2010, Ghosh and Kobayashi 2011, Sadiq, Bostan et al. 2018), and their proficiency in infecting a broad range of mammalian hosts including many domesticated animal species (Martella, Banyai et al. 2010, Doro, Farkas et al. 2015), the high frequencies of hosts infected with multiple genotypes is not entirely surprising. These coinfections present abundant opportunities for rotavirus recombination.

### Rotavirus Recombination Generates Genetic Diversity

Homologous recombination previously has not been considered a significant driver in rotavirus genetic diversity and evolution (Ramig 1997, Woods 2015). Recombination is usually expected to be deleterious as the breakage of open reading frames may disrupt RNA secondary structure and alter protein functionality (Lai 1992, Simon-Loriere and Holmes 2011). However, recombination, as with reassortment (Ramig and Ward 1991, Iturriza-Gomara, Isherwood et al. 2001, Schumann, Hotzel et al. 2009, Ghosh and Kobayashi 2011, Jere, Chaguza et al. 2018), may further increase rotavirus genetic diversity due to epistatic interactions resulting in reassortant-specific or recombinant-specific mutations (Zeldovich, Liu et al. 2015). Formerly deleterious mutations may become beneficial when the genetic background changes, resulting in an increase in circulating pathogenically relevant viral strains.

In our study, most recombination events occurred between strains of the same genotype (Fig. 1; Table 2; Supplementary Table 1). This outcome is consistent with the expectation that intragenotypic is more common since it would less likely to disrupt protein or secondary RNA structure. Nonetheless, intragenotypic recombination can have long lasting effects on rotavirus genetic diversity. For example, we identified a recombinant sub-lineage within the R2 clade of segment 1, the polymerase-encoding segment (Fig. 2; Fig. 3; Table 2). As the same event is found in strains isolated years apart from geographically distant locations, we can infer that the resulting genotype was sufficiently fit enough to be maintained in the population and disperse widely (Supplementary Table 1). This finding suggests that this homologous recombination event has had a long-term effect on rotavirus diversity.

While comparatively less common, we observed instances of intergenotypic recombination in all segments with the exception of segment 7 (NSP3), the only segment where we observed no recombination events (Table 2; Supplementary Table 1). A previous study reported intergenotypic recombination events in segment 6 (VP6), segment 8 (NSP2), and segment 10 (NSP4) (Jere, Mlera et al. 2011), so our study adds to the number of segments able to tolerate intergenotypic recombination. Interestingly, the serotype proteins, VP4 and VP7, have the most different genotypes, with 51 and 36 respectively (Steger, Boudreaux et al. 2019). Given this genetic diversity, the chances of two viruses with different G or P types coinfecting a cell is substantially higher than other segments. In addition, both VP4 and VP7 seem to be more prone to reassortment, and to tolerate more divergent genetic backgrounds or genome constellations (Martella, Ciarlet et al. 2003, Gentsch, Laird et al. 2005, McDonald, Matthijnssens et al. 2009, Patton 2012). This diversity and tolerance of many different genetic backgrounds implies that VP4 and VP7 may be more tolerant of recombination between divergent strains than the other segments. Our data seem to support this claim.

Specifically, we observed numerous instances of intergenotypic recombination in segment 9, the VP7 coding segment (Table 2; Supplementary Table 1). These events appear to be beneficial because the recombinant genotypes persisted in populations long enough for multiple samples showing the same event to be sampled (Table 2). For example, the same mosaic VP7 G6 gene (Fig. 8) was sequenced in multiple bovine strains, one isolated in 2009 (HM591496) and another in 2012 (KF170899), by two separate research groups. While still differentiable, the parents are closely related. However, in some instances, events between highly divergent genotypes seem to have been able to persist in populations. Both a 2014 Malawi isolate (MG181727) and a 2006 isolate from the United States (KC443034) showed a similar G1-G2 VP7 mosaic gene (Fig. 8). The fact that these two G genotypes are highly divergent from one another and were identified in different years in different locations by different research groups supports the contention that it is a true recombination event. Altogether, 22/24 instances of recombination in segment 9 occurred between different G serotypes (Supplementary Table 1). These examples indicate that mosaic genes formed from two divergent genotypes are relatively common and increase the diversity of circulating VP7 genotypes.

In addition, 7/16 instances of recombination in segment 4 (VP4) were intergenotypic. Recombination was observed between P4-P8, P6-P8, and P8-P14 serotypes (Supplementary Table 1). P4, P6 and P8 are all relatively closely related being in the P[II] genogroup, while P14 is in the P[III] genogroup. These putative recombination events suggest that relevant serotype diversity in human hosts is expanding. However, as P4, P6, and P8 are also the dominant P types in human infections, there is a sampling bias towards this genogroup as the genotypes within this group are more likely to coinfect humans, so caution should be taken with this conclusion.

Segment 5 (NSP1) also showed recombination between highly divergent strains. Not only is NSP1 not strictly required for viral replication (Hua, Chen et al. 1994), it is also the least conserved of all rotavirus proteins, including even the serotype proteins VP4 and VP7 (Arnold and Patton 2011), suggesting that intergenotypic recombination disrupting the protein’s amino acid sequence may be less likely to be deleterious.

Segments 7 (NSP3), 8 (NSP2), 10 (NSP4), and 11 (NSP5) all had low rates of recombination. These segments are short thus recombination is expected to be less likely, however segment 9 (VP7), one of the smallest segments, defies this pattern, having a high number of events observed. The smaller segments may also be less able to tolerate recombination events due to the important roles they play during the formation and stabilization of the supramolecular RNA complex (Fajardo, Sung et al. 2015) during rotavirus packaging and assembly (Li, Manktelow et al. 2010, Suzuki 2015, Borodavka, Dykeman et al. 2017, Fajardo, Sung et al. 2017).

### Generation of Escape Mutants

Recombination involving regions encoding conserved epitopes, especially in segments that encode proteins involved in host cell attachment and entry, may provide selective advantages to rotaviruses by allowing them to evade inactivation by host-produced antibodies. These escape mutants may be generally less fit than wildtype viruses, but competitively advantaged in hosts because of a lack of host recognition. Subsequent intrahost adaptation may then select for compensatory fitness-increasing mutations allowing these strains to be competitive with circulating rotavirus strains. Many of the recombination events observed in our study appeared to generate such escape mutants. For example, we found two instances of the same segment 9 (VP7) G1-G3 recombination event (Table 2; KJ751729 and KP752817) with amino acid changes in two of four conserved epitope regions due to the recombination event, which suggests that this strain prevailed in the population because it was better able to evade antibody neutralization.

Based on the I-TASSER structural predictions, the putative recombinant VP7 proteins detected in our survey appear able to fold properly and form functional proteins despite containing amino acid sequence from different ‘parental’ genotypes (Fig. 6). While the secondary structure appeared slightly altered (e.g., shorter beta sheets), the recombinant VP7 proteins generally maintained their 3-dimensional shape. The selective advantage from swapping epitopes may outweigh any potential decrease in protein stability resulting from recombination.

We also identified many recombination events involving segment 4 (VP4). In order to infect cells, VP4 must be proteolytically cleaved to produce VP5* and VP8* (Arias, Romero et al. 1996). Most of the segment 4 recombination events involved the spike head of the VP8* protein or the spike body/stalk region of the VP5* protein (antigen domain). Escape mutant studies (Zhou, Burns et al. 1994, Ludert, Ruiz et al. 2002, Aoki, Settembre et al. 2009, Nair, Feng et al. 2017) for VP4 show the VP8* spike head recognizes histo-blood group antigens, which is one of rotavirus’s main host range expansion barriers (Huang, Xia et al. 2012, Hu, Sankaran et al. 2018, Lee, Dickson et al. 2018). VP5* mediates membrane penetration during cell entry (Yoder and Dormitzer 2006). VP4 recombinants therefore may help the virus expand host range and aid in immune evasion.

Segment 6 (VP6) also showed recombination events resulting in substantial amino acid changes. VP6 is a more conserved protein, but also an antigenic protein that interacts with naïve B cells (Parez, Garbarg-Chenon et al. 2004). This feature suggests that there may be selection for VP6 escape mutants to evade host immune responses. Structure analyses of VP6 indicated that it is relatively conserved across genotypes (Jiang, Tsunemitsu et al. 1992, Tang, Gilbert et al. 1997, Charpilienne, Lepault et al. 2002), which may explain why VP6 seems to have more frequent intergenotypic recombination. Conserved epitopes in VP6 exist around amino acid positions 197 to 214, and 308 to 316 (Aiyegbo, Eli et al. 2014).

### Recombination in Other dsRNA Viruses

Recombination has had a significant impact on the diversity of other dsRNA reoviruses (He, Ding et al. 2010). One study of 692 complete bluetongue virus segments found evidence for at least 11 unique recombinant genotypes (1.6%) (He, Ding et al. 2010). The case for recombination among bluetongue viruses is strengthened by the fact that viruses containing the same (or similar) recombinant segments were isolated by different research groups in different countries at different times, indicating that the recombinant viruses persisted and spread following the recombination event (Carpi, Holmes et al. 2010, He, Ding et al. 2010). Another study found multiple possible instances of recombination in genome segment 8 (encoding NS2) of the epizootic hemorrhagic disease virus, a reovirus similar to bluetongue virus (Anthony, Maan et al. 2009). Several studies have reported recombination among the dsRNA rice black-streaked dwarf virus, which is also a member of the *Reoviridae* family, but infects plants (Li, Xia et al. 2013, Yin, Zheng et al. 2013). Putative recombinants were identified in six of the ten Southern rice black-streaked dwarf virus segments (Li, Xia et al. 2013, Yin, Zheng et al. 2013). Finally, intragenic recombination was observed in multiple isolates of the African horse sickness virus, an *Oribivirus* of the family *Reoviridae* (Ngoveni, van Schalkwyk et al. 2019). At least one of these events appeared in multiple subsequent lineages (Ngoveni, van Schalkwyk et al. 2019).

In study of the family *Birnaviridae*, 1,881 sequences were analyzed for evidence of recombination (Hon, Lam et al. 2008). While no interspecies recombination was observed, at least eight putative instances of intraspecies recombination were observed among the infectious bursal disease viruses and the aquabirnaviruses (Hon, Lam et al. 2008). Subsequent studies focusing on the infectious bursal disease viruses supported these results, and identified additional potential recombination events (He, Ma et al. 2009, Jackwood 2012, Vukea, Willows-Munro et al. 2014). We note that birnaviruses’ genetic material is in a complex with ribonucleoprotein, while the genetic material of *Reoviridae* members is free in the virion, which could be a factor in differing rates of recombination across these dsRNA viruses.

Recombination has also been observed in dsRNA mycoviruses, including in the *Partitiviridae* (Botella, Tuomivirta et al. 2015) and the *Hypoviridae* (Carbone, Liu et al. 2004, Linder-Basso, Dynek et al. 2005, Feau, Dutech et al. 2014) and the *Totiviridae* (Voth, Mairura et al. 2006). Recombination in Gammapartititvirus, which infects the fungus *Gremmeniella abietina*, may have permitted the virus to cross species borders (Botella, Tuomivirta et al. 2015). In cryphonectria hypovirus 1, which infects chestnut blight, recombination was implicated in the spread of the virus in Europe (Feau, Dutech et al. 2014). Collectively, these studies, and those of other dsRNA families, suggest that not only is recombination possible, but also significantly impacts virus evolution. There is little doubt that this conclusion will be strengthened as more dsRNA viruses are discovered and/or sequenced.

### Possible Mechanism of Recombination in Rotavirus

The precise mechanism for how rotavirus recombination occurs is unknown, but inferences can be made because many of the details regarding rotavirus replication and packaging have been resolved (McDonald and Patton 2011, Borodavka, Desselberger et al. 2018). The most-accepted hypothesis is that recombination takes place when the rotavirus +ssRNA is replicated after being packaged in the nucleocapsid (Esona, Roy et al. 2017, Jing, Zhang et al. 2018). For packaging and replication to occur, the eleven +ssRNA segments must join a protein complex consisting of the VP1 polymerase and the VP3 capping enzyme. Secondary RNA structures in the non-translated terminal regions (NTRs) aid in the formation of this supramolecular RNA complex (Fajardo, Sung et al. 2015, Borodavka, Dykeman et al. 2017) and determine whether the segments are packaged (Li, Manktelow et al. 2010, Suzuki 2015). Perhaps recombination occurs when multiple homologous RNA strands are joined in the same complex, allowing the homologous NTRs to partially hybridize. In this scenario, the VP1 polymerase replicates part of one strand before switching to the other, thus producing a recombined segment. This conjecture is supported by the fact that recombination tends to occur in segment regions where self-hybridization forms three-dimensional structures. Moreover, rotavirus do not seem constrained from packaging extra genetic material (Desselberger 1996). A similar form of template switching is seen in poliovirus, an ssRNA virus, which exhibits high rates of recombination, although the precise mechanism may be different in that case insofar as in poliovirus, the polymerase may be stalled due to a hairpin or other secondary structure, and switches to a different template (Tolskaya, Romanova et al. 1987). Further study is needed to determine the precise mechanism of recombination in rotavirus.

## SUPPLEMENTARY INFORMATION

**Supplementary Table 1.** Putative recombination events identified by RDP4 for whole genome Rotavirus A sequences. A total of 117 events were identified by at least 6/7 RDP4 programs. Shaded rows indicate strongly supported events (identified by 7/7 RDP4 programs or found in multiple sequenced isolates). Events that were flagged by RDP4, but were not strongly supported are not listed here.

| Segment | Accession Number | Breakpoint Cis (Excluding Gaps) | p-Values | Representative Major Parent Sequence, Source Species, Year of Isolation, % ANI, Genogroup | Representative Minor Parent Sequence, Source Species, Year of Isolation, % ANI, Genogroup |
| --- | --- | --- | --- | --- | --- |
| VP1 | JQ988899, *Homo sapiens*, Croatia, 2006 | 970 (781-986) - 1734 (1693-1751) | RDP 1.02E-43<br>GENECONV. 8.72E-48<br>BootScan 4.17E-32<br>MaxChi. 1.15E-17<br>Chimera 1.05E-17<br>SiScan 4.30E-19<br>3Seq 2.96E-09 | KX632342, *Homo sapiens*, Uganda, 93.7%, R1 | GQ414540, *Homo sapiens*, Germany, 2009, 98.7%, R2 |
| VP1 | KU739903, *Sus scrofa*, Taiwan, 2015 | 1-(747-805) | RDP 1.19E-24<br>GENECONV 1.41E-21<br>BootScan. 5.14E-21<br>MaxChi. 8.41E-16<br>Chimera 3.57E-16<br>SiScan. 9.64E-16<br>3Seq 2.83E-09 | KU739900, *Sus scrofa*, Taiwan, 2015, 95.8%, R1 | KU739904, *Sus scrofa*, Taiwan, 2015, 99.1%, R1 |
| VP1 | KF812721, *Homo sapiens*, South Korea, 2010 | 2504 (2464-2537) - 3268 (3232-116) | RDP 2.58E-22<br>GENECONV 2.12E-08<br>BootScan 4.71E-22<br>MaxChi 9.96E-16<br>Chimera 1.47E-13<br>SiScan 5.10E-09<br>3Seq 2.05E-02 | JX027681, *Homo sapiens*, Australia, 2007, 98.1%, R1 | JF796734, *Sus scrofa*, South Korea, 2006, 94.6%, R1 |
| VP1 | JQ069926, *Homo sapiens*, Canada, 2008 | 2498 (2356-2507) - 3268 | RDP 9.14E-22<br>GENECONV 5.71E-19<br>BootScan 1.00E-18<br>MaxChi 1.30E-15<br>Chimera 2.51E-15<br>SiScan. 7.42E-23<br>3Seq 2.78E-25 | JX195074, *Homo sapiens*, Italy, 2010, 98.1%, R1 | JX027681, *Homo sapiens*, Australia, 2007, 99.6%, R1 |
| VP1 | KM454503, *Equus caballus*, USA, 1981 | 840 (802-951) - 3281 | RDP 2.77E-15<br>GENECONV 1.36E-19<br>BootScan 4.79 E-21<br>MaxChi 1.55E-17<br>SiScan 1.27E-40<br>3Seq 8.22E-05 | Unknown, closest = DQ838638, unknown, R2 | KM454492, *Equus caballus*, United Kingdom, 1976, 99.1%, R2 |
| VP1 | KU714444, *Homo sapiens*, Malawi, 2000 | 1656 (1636-1673) - 1752 (1740-1766) | RDP 1.35E-18<br>GENECONV 1.25E-17<br>BootScan 3.76E-11<br>MaxChi 3.81E-02<br>Chimera 4.33E-02<br>3Seq 5.67E-09 | JN706454, *Homo sapiens*, Thailand, 2009, 97.4%, R1 | KX655451, *Homo sapiens*, Uganda, 2013, 97.9%, R2 |
| VP1 | JQ069936, *Homo sapiens*, Canada, 2009 | 1670 (1531-1703) - 2419 (2308-2436) | RDP 1.83E-14<br>GENECONV 7.68E-13<br>MaxChi 2.73E-08 | KU248405, *Homo sapiens*, Bangladesh, 2010, 98.7%, R2 | HQ657171, *Homo sapiens*, South Africa, 2009, 99.7%, R2 |
|  |  |  | Chimera 2.48E-08<br>SiScan 2.65E-08<br>3Seq 5.15E-19 |  |  |
| VP1 | KU739903, *Sus scrofa*, Taiwan, 2015 | 1543 (138-1588) - 2220 (2160-2260) | RDP 3.06E-13<br>GENECONV 6.19E-04<br>BootScan 9.40E-11<br>MaxChi 2.98E-08<br>Chimera 5.67E-08<br>SiScan 1.19 E-06<br>3Seq 2.83E-09 | KU739900, *Sus scrofa*, Taiwan, 2015, 97.8%, R1 | KU739904, *Sus scrofa*, Taiwan, 2015, 96.5%, R1 |
| VP1 | JQ988899, *Homo sapiens*, Croatia, 2006 | 222 (3094-362) - 969 (882-NA) | RDP 3.76E-12<br>GENECONV 4.88E-10<br>MaxChi 7.74E-09<br>Chimera 4.5E-07<br>SiScan 6.32E-14<br>3Seq 5.64E-09 | KJ559215, *Homo sapiens*, Argentina, 1998, 96.8%, R1 | Unknown, closest = JX195074, *Homo sapiens*, Italy, 2010, R1 |
| VP1 | KC580176, *Homo sapiens*, USA, 1988 | 1471 (1328-1521) - 3270 | RDP 9.77E-12<br>GENECONV 3.81E-08<br>MaxChi 6.16E-12<br>Chimera 2.36E-04<br>SiScan 7.42E-23<br>3Seq 1.91E-20 | KC580526, *Homo sapiens*, USA, 1979, 99%, R1 | KC580601, *Homo sapiens*, USA, 1988, 100%, R1 |
| VP1 | JQ069918, *Homo sapiens*, Canada, 2008 | 604 (541-662) - 956 (889-1068) | RDP 2.42E-10<br>GENECONV 3.25E-08<br>BootScan 2.38E-09<br>MaxChi 5.62E-03<br>Chimera 5.62E-03<br>SiScan 1.10E-04<br>3Seq 1.13E-08 | KJ751834, *Homo sapiens*, South Africa, 2009, 99.5%, R1 | JQ069924, *Homo sapiens*, Canada, 2008, 99.7%, R1 |
| VP1 | KC579647, *Homo sapiens*, USA, 1988 | 35 – 1448 (1290-1498) | GENECONV 4.30E-04<br>BootScan 2.63E-05<br>MaxChi 2.34E-09<br>Chimera 1.41E-08<br>SiScan 6.55E-11<br>3Seq 4.34E-19 | KC580601, *Homo sapiens*, USA, 1988, 100%, R1 | KC579509, *Homo sapiens*, USA, 1989, 99.4%, R1 |
| VP1 | KU199270, *Homo sapiens*, Bangladesh, 2010 | 670 (541-711) - 1403 (1327-1445) | Flagged for 285 sequences, p-value = 3.33E-09 | KX536654, *Homo sapiens*, India, 2011, 98.6%, R2 | KU356640, *Homo sapiens*, Bangladesh, 2013, 100%, R2 |
| VP1 | KU356662, KU356640, *Homo sapiens*, 2013 | 656 (541-711) - 2661 (2600-2735) | RDP 5.87E-08<br>GENECONV 1.43E-04<br>MaxChi 9.02E-10<br>SiScan 6.71E-20<br>3Seq 6.88 E-23 | KU356607, *Homo sapiens*, Bangladesh, 2013, 99.5%, R2 | KC178768, *Homo sapiens*, Italy, 2007, 99.6%, R2 |
| VP1 | KU199270, *Homo sapiens*, Bangladesh, 2010 | 3282 (3116-50) - 1410 (1363-1478) | GENECONV 3.94E-06<br>BootScan 2.03E-04<br>MaxChi 1.20E-09<br>Chimera 3.36E-10<br>SiScan 2.57E-09<br>3Seq 5.81E-18 | JQ069920, *Homo sapiens*, Canada, 2008, 99.5%, R2 | KJ751889, *Homo sapiens*, Ethiopia, 2009, 99.1%, R2 |
| VP2 | KJ753425, *Homo sapiens*, Uganda, 2011 | 599 (561-623) - 991 (967-1019) | RDP 6.28E-47<br>GENECONV 5.13E-45<br>BootScan 2.53E-44<br>MaxChi 5.39E-12<br>Chimera 6.10E-12<br>SiScan 8.38E-15<br>3Seq 1.96E-09 | JF490148, *Homo sapiens*, Australia, 2004, 98.3%, C1 | KP752972, *Homo sapiens*, Gambia, 2008, 99.7%, C2 |
| VP2 | KU714445, *Homo sapiens*, Malawi, 2000 | 1155(1139-1170) - 1887 (1862-1908) | RDP 2.37E-41<br>GENECONV 1.62E-33<br>BootScan 5.28E-39<br>MaxChi 1.26E-16<br>Chimera 1.42E-16<br>SiScan 8.61E-25<br>3Seq 1.96E-09 | KP752983, *Homo sapiens*, South Africa, 2009, 94.3%, C1 | KU714456, *Homo sapiens*, Malawi, 2000, 99.9%, C2 |
| VP2 | KX655439, *Homo sapiens*, Uganda, 2013 | 2713 (2672-2717) - 158 (143-171) | RDP 1.34E-25<br>GENECONV 9.67E-24<br>MaxChi 1.54E-03<br>Chimera 8.56E-04<br>SiScan 1.61E-05<br>3S— 2.15E--08 | Unknown, closest = KX632343, *Homo sapiens*, Uganda, 2013, C2 | MG181349, *Homo sapiens*, Malawi, 2002, 99.4%, C1 |
| VP2 | KJ753617, *Homo sapiens*, South Africa, 2004 | 548 (464-560) - 784 (754-842) | RDP 5.31E-27<br>GENECONV 4.03E-24<br>BootScan 7.13E-23 | KJ752329, *Homo sapiens*, South Africa, 2003, 99.4%, C1 | KX655439, *Homo sapiens*, Uganda, 2013, 98.4%, C2 |
|  |  |  | MaxChi 3.85E-05<br>Chimera 4.51E-05<br>3Seq 1.96E-09 |  |  |
| VP2 | HM988959, *Bos taurus*,<br>South Korea, Unknown | 558 (524-586) -<br>2685 | RDP 1.14E-21<br>GENECONV 2.68E-25<br>BootScan 2.00 E-21<br>MaxChi 1.29E-05<br>Chimera 1.11E-03<br>SiScan 3.88E-15<br>3Seq 5.94E-21 | JQ069842, *Homo sapiens*, Canada,<br>2008, 95.2%, C1 | JX971570, *Bos taurus*,<br>South Korea, 2004, 100%,<br>C1 |
| VP2 | MG181591, *Homo sapiens*,<br>Malawi, 2013 | 1129 (1106-1140) -<br>1299 (1292-1318) | RDP 3.47 E-21<br>GENECONV 2.01E-19<br>BootScan 4.15E-15<br>MaxChi 4.04E-03<br>Chimera 3.95E-03<br>3Seq 1.96E-09 | KJ753828, *Homo sapiens*, Zimbabwe,<br>unknown, 99.1%, C2 | KJ753425, *Homo sapiens*,<br>Uganda, 2011, 99.4%, C1 |
| VP2 | KX632343, *Homo sapiens*,<br>Uganda, 2013 | 2666 (2614-39) -<br>593 (560-612) | RDP 9.89E-21<br>GENECONV 2.26 E-17<br>BootScan 5.20E-19<br>MaxChi 4.65E-13<br>Chimera 3.18 E-13<br>SiSeq 2.53E-24<br>3Seq 1.37E-05 | JN258812, *Homo sapiens*, Belgium,<br>2000, 91.7%, C1 | KJ753402, *Homo sapiens*,<br>South Africa, 2005,<br>99.5%, C2 |
| VP2 | KX632343, *Homo sapiens*,<br>Uganda, 2013 | 1357 (1322-1368) -<br>1808 (1790-1830) | RDP 1.14E-21<br>GENECONV 3.44E-17<br>MaxChi 1.69E-11<br>Chimera 3.57E-10<br>SiScan 2.84E-14<br>3Seq 1.96E-09 | MG181371, *Homo sapiens*, Malawi,<br>2002, 95.3%, C1 | KJ870879, *Homo sapiens*,<br>Democratic Republic of<br>the Congo, 99.3%, C2 |
| VP2 | KY055428, *Capra aegagrus*,<br>Uganda, 2014 | 983 (960-1009) -<br>1238 (1194-1243) | RDP 1.64E-18<br>GENECONV 1.12E-16<br>BootScan 2.13E-10<br>MaxChi 1.67E-05<br>Chimera 4.73E-04<br>SiScan 4.35E-05<br>3Seq 1.96E-09 | JN831232, *Bos taurus*, South Africa,<br>2007, 94.7%, C2 | KF636257, *Bos taurus*,<br>South Africa, 2007,<br>99.6%, C2 |
| VP2 | KJ870901, *Homo sapiens*,<br>Democratic Republic of the<br>Congo, Unknown | 2330 (2304-2359) -<br>2648 (2589-1) | RDP 1.96E-18<br>GENECONV 2.99E-16<br>BootScan 5.51E-16<br>MaxChi 1.71E-03<br>Chimera 7.67E-04<br>SiScan 6.29E-06<br>3Seq 1.96E-09 | KY055417, *Sus scrofa*, Uganda, 2016,<br>95%, C1 | LC367260, *Homo sapiens*, Nepal, 2008,<br>99.4%, C1 |
| VP2 | KC257092, *Camelus dromedaries*,<br>Sudan, 2002 | 2047 (2017-2102) -<br>733 (661-754) | RDP 2.53E-16<br>GENECONV 4.72E-06<br>BootScan 7.95E-23<br>MaxChi 2.68E-13<br>Chimera 8.40E-12<br>SiScan 8.51E-11<br>3Seq 9.20E-08 | Unknown, closest =<br>DQ480724, *Homo sapiens*, Iran,<br>unknown, C2 | FJ347123, *Lama guanicoe*, Argentina,<br>1998, 99.9%, C2 |
| VP2 | KX632343, *Homo sapiens*,<br>Uganda, 2013 | 1962 (1932-1978) -<br>2133 (2099-2136) | RDP 1.44E-13<br>GENECONV 7.46E-10<br>MaxChi 3.26E-06<br>Chimera 4.13E-05<br>SiScan 1.49E-07<br>3Seq 1.96E-09 | JN706477, *Homo sapiens*, Thailand,<br>2010, 93%, C1 | KX655439, *Homo sapiens*, Uganda, 2013,<br>97.7%, C2 |
| VP2 | KJ627064, *Homo sapiens*,<br>Paraguay, 2002 | 2675 (2615-95) -<br>409 (370-449) | RDP 7.01E-06<br>GENECONV 2.53E-03<br>BootScan 1.26E-04<br>MaxChi 4.49E-02<br>Chimera 3.89E-02<br>SiScan 1.20E-02<br>3Seq 9.21E-04 | KJ752074, *Homo sapiens*, South Africa,<br>2005, 98.5%, C1 | KJ752859, *Homo sapiens*,<br>Zimbabwe, 2011, 100%,<br>C1 |
| VP3 | KJ412679, *Homo sapiens*,<br>Paraguay, 2009 | 1 (2497-3) - 1068<br>(1047-1080) | RDP 1.04E-72<br>GENECONV 1.11E-59<br>BootScan 1.41E-73<br>MaxChi 3.58E-31<br>Chimera 5.31E-31<br>SiScan 6.77E-38<br>3Seq 2.83E-09 | JX185760, *Homo sapiens*, Italy, 2007,<br>97.8%, M1 | KJ412622, *Homo sapiens*,<br>Paraguay, 2009, 99.9%,<br>M3 |
| VP3 | KP007164, *Homo sapiens*, Philippines, 2012 | 968 (968-996) - 1753 (1727-1777) | RDP 1.97E-38<br>GENECONV 2.93E-35<br>BootScan 2.16E-37<br>MaxChi 2.45E-16<br>Chimera 1.76E-15<br>SiScan 2.85E-19<br>3Seq 2.83E-19 | KY000551, *Homo sapiens*, Germany, 2016, 99.4%, M2 | KP882081, *Homo sapiens*, Bangladesh, 2007, 99.7%, M2 |
| VP3 | KX171542, *Homo sapiens*, South Korea, 2014 | 1465 (1432-1538) - 2482 (2461-59) | RDP 1.40E-37<br>GENECONV 7.17E-32<br>BootScan 2.84E-35<br>MaxChi 2.03E-19<br>Chimera 3.57E-19<br>SiScan 3.27E-23<br>3Seq 2.83E-09 | KJ752827, *Homo sapiens*, South Africa, 2003, 98.8%, M2 | KX171538, *Homo sapiens*, South Korea, 2012, 99.7%, M2 |
| VP3 | KX171543, *Homo sapiens*, South Korea, 2014 | 1650 (1607-1688) - 2472 (2455-59) | RDP 5.19E-37<br>GENECONV 2.46E-33<br>BootScan 2.81E-34<br>MaxChi 1.95E-17<br>Chimera 1.05E-17<br>SiScan 1.55E-19<br>3Seq 2.83E-09 | KU248374, *Homo sapiens*, Bangladesh, 2010, 99.1%, M2 | KC443523, *Homo sapiens*, Australia, 2007, 99.6%, M2 |
| VP3 | AB779630, *Sus scrofa*, Thailand, 2008 | 1590 (1578-1645) - 90 (2490-96) | RDP 3.19E-33<br>GENECONV 1.69E-25<br>BootScan 2.39E-31<br>MaxChi 2.05E-16<br>Chimera 3.42E-15<br>SiScan 9.38E-25<br>3Seq 1.42E-24 | MG781054, *Sus scrofa*, 2010-2011, Thailand, 92.7%, M1 | AB779632, *Sus scrofa*, Thailand, 2009, 99.9%, M1 |
| VP3 | MG181592, *Homo sapiens*, Malawi, 2013 | 2334 (2318-2349) - 196 (150-204) | RDP 1.09E-32<br>GENECONV 6.85E-18<br>BootScan 5.72E-27<br>MaxChi 6.16E-12<br>Chimera 9.20E-12<br>SiScan 3.92E-10<br>3Seq 3.54E-27 | KC442894, *Homo sapiens*, USA, 2008, 96.4%, M2 | KT919912, *Homo sapiens*, USA, 2013, 99%, M1 |
| VP3 | JN013991, *Homo sapiens*, South Africa, 2008 | 82 (2550-99) - 260 (253-268) | RDP 1.69E-29<br>GENECONV 8.98 E-27<br>BootScan 1.24E-17<br>MaxChi 9.47E-06<br>Chimera 1.58E-05<br>SiScan 1.76E-02<br>3Seq 5.67E-09 | KJ752816, *Homo sapiens*, South Africa, 2011, 97.4%, M1 | KP883027, *Homo sapiens*, Mali, 2008, 99.4%, M2 |
| VP3 | KU356576, *Homo sapiens*, Bangladesh, 2010 | 844 (815-855) - 1194 (1142-1219) | RDP 4.55E-24<br>GENECONV 1.69E-21<br>BootScan 3.82E-20<br>MaxChi 1.89E-09<br>Chimera 1.77E-09<br>SiSeq 1.18E-07<br>3Seq 2.83E-09 | KC178787, *Homo sapiens*, Italy, 2008, 97.4%, M2 | KP882081, *Homo sapiens*, Bangladesh, 2007, 100%, M2 |
| VP3 | JQ0692727, *Homo sapiens*, Canada, 2007 | 51 (2482-59) - 845 (817-932) | RDP 4.27E-23<br>GENECONV 9.70E-19<br>BootScan 6.96E-18<br>MaxChi 8.76E-14<br>Chimera 4.71E-14<br>SiScan 2.58E-17<br>3Seq 3.98E-35 | JQ069748, *Homo sapiens*, Canada, 2008, 99.1%, M1 | JQ069762, *Homo sapiens*, Canada, 2008, 99.7%, M1 |
| VP3 | JQ069754, *Homo sapiens*, Canada, 2008 | 835 (811-879) - 1621 (1591-1631) | RDP 6.24E-20<br>GENECONV 4.13E-17<br>BootScan 4.05E-17<br>MaxChi 1.69E-09<br>Chimera 1.31E-09<br>SiScan 1.91E-10<br>3Seq 1.71E-27 | KJ627117, *Homo sapiens*, Paraguay, 1999, 99%, M1 | KP645258, *Homo sapiens*, Australia, 2010, 99.7%, M1 |
| VP3 | MG181592, *Homo sapiens*, Malawi, 2013 | 1736 (1712-1755) - 1941 (1889-1953) | RDP 2.97E-19<br>GENECONV 7.80E-18<br>BootScan 5.65E-15<br>MaxChi 5.47E-04<br>Chimera 5.27E-04<br>SiSeq 9.43E-07<br>3Seq 2.83E-09 | KC178787, *Homo sapiens*, Italy, 2008, 97%, M2 | LC374087, *Homo sapiens*, Nepal, 2009, 100%, M1 |
| VP3 | KJ753665, *Homo sapiens*, South Africa, 2004 | 2158 (2127-2201) - 2517 (2447-64) | Flagged in 107, KA p-value= 2.23E-18, Global p-value 5.69E-11 | KJ753481, *Homo sapiens*, Ethiopia, 2012, 94.9%, M1 | KY055418, *Sus scrofa*, Uganda, 2016, 98.6%, M1 |
| VP3 | AB374145, *Bos taurus*, Japan, Unknown | 810 (765-875) - 1544 (1486-1639) | RDP 1.85E-08<br>GENECONV 1.83E-04<br>BootScan 3.64E-05<br>MaxChi 1.68E-06<br>Chimera 3.72E-06<br>SiScan 2.14E-07<br>3Seq 4.41E-10 | LC340015, *Homo sapiens*, Japan, 2014, 95.5%, M2 | KJ411434, *Homo sapiens*, USA, 2012, 97.4%, M2 |
| VP3 | KJ753665, *Homo sapiens*, South Africa, 2004 | 1212 (1140-1312) - 1732 (1632-1801) | RDP 3.84E-07<br>GENECONV 3.50E-05<br>BootScan 1.92E-05<br>MaxChi 2.67E-05<br>Chimera 1.59E-05<br>SiScan 3.24E-09<br>3Seq 2.08E-04 | HQ392255, *Homo sapiens*, Belgium, 2008, 96.2%, M1 | KX778614, *Homo sapiens*, Dominican Republic of the Congo, 2013, 99.4%, M1 |
| VP3 | KJ753665, *Homo sapiens*, South Africa, 2004 | 580 (498-612) - 922 (831-964) | GENECONV 7.32E-11<br>BootScan 1.64E-09<br>MaxChi 3.90E-05<br>Chimera 3.85E-06<br>SiScan 9.81E-09<br>3Seq 7.82E-20 | KP752825, *Homo sapiens*, South Africa, 2002, 99.6%, M1 | JQ069760, *Homo sapiens*, Canada, 2008, 99.7%, M1 |
| VP3 | KX655453, *Homo sapiens*, Uganda, 2013 | 1258 (872-1389) - 1798 (1662-1917) | Flagged in 551 sequences, Global p-value= 1.78E-12 | KP882576, *Homo sapiens*, Ghana, 2009, 94.9%, M2 | KU714446, *Homo sapiens*, Malawi, 2000, 96.7%, M2 |
| VP4 | KX655509, *Homo sapiens*, Uganda, 2013 | 321 (308-323) - 683 (616-634)<br>VP8* domain | RDP 2.16E-54<br>GENECONV 1.17E052<br>BootScan 1.59E-51<br>MaxChi 7.47E-19<br>Chimera 1.44E-18<br>SiScan 8.23E-23<br>3Seq 2.07E-09 | KP902549, *Homo sapiens*, South Africa, 2003, 99.3%, P8 | KJ52687, *Homo sapiens*, South Africa, 2000, 99.3%, P8 |
| VP4 | KX655487, *Homo sapiens*, Uganda, 2013 | 334(326-347) - 682(670-698)<br>VP8* domain | RDP 1.06E-55<br>GENECONV 5.94E-51<br>BootScan 1.77E-50<br>MaxChi 4.55E-19<br>Chimera 5.96E-19<br>SiScan 1.60E-17<br>3Seq 2.07E-09 | KX646644, *Homo sapiens*, India, 2012, 96.6%, P6 | MG181340, *Homo sapiens*, Malawi, 2002, 99.7%, P8 |
| VP4 | MG181725, *Homo sapiens*, Malawi, 2014 | 374 (367-382) - 542 (526-553)<br>VP8* domain | GENECONV 5.45E-46<br>BootScan 2.45E-41<br>MaxChi 4.02E-11<br>Chimera 3.97E-11<br>SiScan 2.83E-13<br>3Seq 2.07E-09 | MG181758, *Homo sapiens*, Malawi, 2014, 99.2%, P8 | MG181890, *Homo sapiens*, Malawi, 2012, 100%, P4 |
| VP4 | KU714447, *Homo sapiens*, Malawi, 2000 | 1380 (1378-1381) - 1562 (1543-1571) | RDP 1.47E-39<br>GENECONV 4.07E-38<br>BootScan 104E-29<br>MaxChi 1.24E-07<br>Chimera 3.25E-07<br>SiScan 7.45E-12<br>3Seq 4.14E-09 | GQ869840, *Homo sapiens*, Malawi, 2000, P8 | KU714458, *Homo sapiens*, Malawi, 2000, 99.5%, P14 |
| VP4 | KX655441, *Homo sapiens*, Uganda, 2013 | 374 (357-382)-484 (475-488)<br>VP8* domain | RDP 5.41E-34<br>GENECONV 1.32E-31<br>BootScan 2.38E-30<br>MaxChi 1.11E-05<br>Chimera 6.90E-06<br>SiScan 4.86E-10<br>3Seq 2.07E-09 | LC260226, *Homo sapiens*, 2015-2016, Indonesia, 99.2%, P6 | JQ069692, *Homo sapiens*, Canada, 2009, 99.1%, P8 |
| VP4 | MG181637, *Homo sapiens*, Malawi, 2013 | 350 (325-368) - 459 (464-466)<br>VP8* domain | GENECONV 5.13E-28<br>BootScan 5.75E-15<br>MaxChi 2.13E-04<br>Chimera 1.00E-04<br>SiScan 4.71E-05<br>3Seq 4.14E-09 | MG181758, *Homo sapiens*, Malawi, 2014, 99.5%, P8 | KJ870892, *Homo sapiens*, Democratic Republic of the Congo, 2007-2010, 95.5%, P6 |
| VP4 | JQ069674, *Homo sapiens*, Canada, 2008 | 1568 (1541-1574) - 2329 (2251-101) | RDP 1.59E-23<br>GENECONV 2.53E-21<br>BootScan 1.59E-23<br>MaxChi 2.96E-12<br>Chimera 2.13E-12<br>SiScan 7.86E-14 | HQ392253, *Homo sapiens*, Belgium, 2008, 99.5%, P8 | HM773714, *Homo sapiens*, USA, 2008, 99.5%, P8 |
|  |  |  | 3Seq 1.44E-35 |  |  |
| VP4 | KC579762, *Homo sapiens*, USA, 1991 | 1073 (1039-1124) - 2332 (2269-8) | RDP 4.67E-23<br>GENECONV 4.72E-17<br>BootScan 8.46E-23<br>MaxChi 1.99E-14<br>Chimera 1.76E-11<br>SiScan 7.07E-25<br>3Seq 4.04E-40 | HQ392253, *Homo sapiens*, Belgium, 2008, 98.7%, P8 | FJ947895, *Homo sapiens*, USA, 1991, 99.9%, P8 |
| VP4 | MG181659, *Homo sapiens*, Malawi, 2013 | 1116 (1064-1146) - 1462 (1402-1501) | RDP 6.53E-11<br>GENECONV 2.20E-09<br>BootScan 1.08E-09<br>MaxChi 1.781E-05<br>Chimera 1.74E-05<br>SiScan 5.94E-06 | MG181725, *Homo sapiens*, Malawi, 2013, 98.1%, P8 | KJ752208, *Homo sapiens*, South Africa, 2012, 98.3%, P4 |
| VP4 | JF490554, *Homo sapiens*, Australia, 2005 | 462 (397-482) - 683 (637-722) | RDP 2.17E-10<br>GENECONV 8.75E-09<br>BootScan 1.89E-10<br>MaxChi 3.26E-04<br>Chimera 1.04E-03<br>SiScan 1.14E-04<br>3Seq 4.14E-09 | KJ751716, *Homo sapiens*, Gambia, 2010, 98.4%, P8 | JQ069674, *Homo sapiens*, Canada, 2008, 99.5%, P8 |
| VP4 | AB008291, *Homo sapiens*, 1995 | 2291 (2223-71) - 807 (762-849) | RDP 1.02E-03<br>BootScan 4.42E-02<br>MaxChi 5.00E-05<br>Chimera 1.32E-04<br>SiScan 1.51E-05<br>3Seq 1.66E-08 | AB039939, *Homo sapiens*, Japan, 1990, 98.6%, P8 | AB008277, *Homo sapiens*, China, 1992, 98.9%, P8 |
| VP6 | KJ482497, *Sus scrofa*, Brazil, 2013 | 589 (584-605) - 801 (790 – 809) | RDP 1.81E-25<br>GENECOV 7.17E-23<br>BootsCan 7.09E018<br>MaxChi 2.98E-10<br>SiScan 2.41E-11<br>3Seq 1.41E-09 | DQ119822, *Sus scrofa*, China, unknown, 98.1%, I2 | KJ482491, *Sus scrofa*, Brazil, 2013, 100%, I5 |
| VP6 | KX632346, *Homo sapiens*, Uganda, 2013 | 1269 (1250-1272) - 56 (1333-103) | RDP 7.77E-19<br>GENECONV 1.33E-17<br>BootsCan 7.27E-09<br>MaxChi 2.1E-02<br>Chimera 3.24E-03<br>SiScan 1.02E-07<br>3Seq 3.06E-11 | KU714448, *Homo sapiens*, Malawi, 2000, 92.7%, I1 | KX655455, *Homo sapiens*, Uganda, 2013, 96.5%, I2 |
| VP6 | JN014005, *Homo sapiens*, South Africa, 2008 | 160 (33-220 - 384 (379-451) | RDP 1.98E-18<br>GENECONV 6.96E-18<br>BootScan 4.63E-16<br>MaxChi 1.00E-06<br>Chimera 2.77E-07<br>3Seq 1.41E-09 | JN706549, *Homo sapiens*, Thailand, 2010, 97.8%, I1 | KJ752622, *Homo sapiens*, Senegal, 2009, 96.4%, I2 |
| VP6 | JX040423, *Homo sapiens*, India, 2009 | 1177(1143-1181) - 160(140-176) | RDP 1.30E-15<br>GENECONV 4.97E-09<br>BootScan 1.95E-11<br>MaxChi 4.38E-11<br>Chimera 9.18E-12<br>SiScan 7.20E-15<br>3Seq 2.82E-09 | KF740531, *Homo sapiens*, India, 2009, I12 | Unknown, closest = KC579918, *Homo sapiens*, USA, 1976), I1 |
| VP6 | JN014003, *Homo sapiens*, South Africa, 2008 | 202 (1333-208) - 384(354-421) | RDP 6.30E-15<br>GENECONV 2.39E-15<br>BootScan 1.04E-12<br>MaxChi 3.76E-05<br>Chimera 3.02E-05<br>SiScan 4.51E-05<br>3Seq 1.41E-09 | KP752725, *Homo sapiens*, Swaziland, 2010, 95.5%, I2 | JX027952, *Homo sapiens*, Australia, 2010, 96.7%, I1 |
| VP6 | KU714448, *Homo sapiens*, Malawi, 2000 | 653(635- 664) - 831(818-846) | RDP 1.24E-12<br>GENECONV 7.55E-10<br>BootScan 5.50E-07<br>MaxChi 1.15E-05<br>Chimera 3.38E-05<br>SiScan 1.38E-04<br>3Seq 1.41E-09 | JN014005, *Homo sapiens*, South Africa, 2008, 95.9%, I1 | KX655455, *Homo sapiens*, Uganda, 2013, 99.4%, I2 |
| VP6 | KU714448, *Homo sapiens*, Malawi, 2000 | 317 (290-333) - 491 (474-507) | RDP 3.94E-15<br>GENECONV 9.37E-14<br>BootScan 2.42E-13<br>MaxChi 2.80E<br>Chimera 2.58E-05 | KX638567, *Homo sapiens*, India, 2010, 96.6%, I1 | GU984758, *Bos taurus*, India, 2007, 98.9%, I2 |
|  |  |  | SiScan 1.73E-07<br>3Seq 1.41E-09 |  |  |
| VP6 | HQ11009, *Homo sapiens*, Belgium, 2007, 98.5% | 754 (697-775) - 1357 (1329-32) | GENECONV 2.23E-05<br>BootScan 1.61E-03<br>MaxChi 3.32E-11<br>Chimera 1.08E-10<br>SiScan 5.16E-14<br>3Seq 2.54E-19 | MF469405, *Homo sapiens*, USA, 2015, 99.3%, I1 | HQ392161, *Homo sapiens*, Belgium, 2007, 98.5%, I1 |
| VP6 | FJ685614, *Homo sapiens*, India, 1992 | 698(672-712) - 1357 (1231-70) | RDP 1.72E-08<br>GENECONV 1.61E-07<br>BootScan 2.85E-08<br>MaxChi 1.40E-09<br>Chimera 2.14E-09<br>SiScan 4.04E-11<br>3Seq 1.54E-23 | KX638599, *Homo sapiens*, India, 2010, 99.4%, I1 | KX632291, *Homo sapiens*, Uganda, 2013, 99.1%, I1 |
| VP6 | KJ753780, *Homo sapiens*, South Africa, 2004 | 513 (463-560) - 1008 (952-1031) | GENECONV 1.89E-05<br>BootScan 2.60E-05<br>MaxChi 9.01E-09<br>Chimera 1.46E-08<br>SiScan 1.38E-09<br>3Seq 1.30E-19 | JN258839, *Homo sapiens*, Belgium, 2001, 99.6%, I1 | LC056797, *Homo sapiens*, Vietnam, 2007, 98.8%, I1 |
| VP6 | HM988972, *Bos taurus*, South Korea, Unknown | 751 (340-811) - 1113 (1080-1132) | RDP 2.08E-08<br>GENECONV 1.92E-08<br>BootScan 2.91E-08<br>MaxChi 1.45E-05<br>Chimera 2.41E-05<br>SiScan 3.56E-05<br>3Seq 5.55E-05 | Unknown, closest = MH238302, *Sus scrofa*, Spain, 2017, I5 | MH423866, *Sus scrofa*, China, 2016, 99.4%, I5 |
| VP7 | AB158431, *Bos taurus*, Japan, 2000 | 901 (868-907)- 1051 (1037-80) | RDP 3.71E-16<br>GENECONV 1.25E-13<br>MaxChi 6.99E-07<br>Chimera 3.69E-05<br>SiScan 8.07E-10<br>3Seq 4.95E-10 | AB077053, *Bos taurus*, Japan, 1995-1996, 95.9%, G8 | U50332, *Bos taurus*, USA, 1991, 97.2%, G6 |
| VP7 | HM591496, *Bos taurus*, India, 2009<br>KF170899, *Bos taurus*, India, 2012 | 1048 (1032-128) - 481 (448-506) | RDP 2.78E-05<br>GENECONV 4.77 E-04<br>BootScan 4.86E-04<br>MaxChi 1.02E-03<br>Chimera 3.78E-05<br>SiSeq 7.21E-09<br>3Seq 3.90E-08 | EF199501, *Bos taurus*, India, 2001-2005, 96.6%, G6 | JX442784, *Bos taurus*, India, 2010, 100%, G6 |
| VP7 | D86274, *Homo sapiens*, 1990 | 353 (338-368) - 745 (730-756) | RDP 1.73E-18<br>GENECONV 1.43E-16<br>BootScan 2.50E-14<br>MaxChi 3.26E-13<br>Chimera 6.54E-13<br>SiScan 2.47E-10<br>3Seq 3.55E-33 | D86282, *Homo sapiens*, Japan, 1981, 98.7%, G3 | KF636217, *Homo sapiens*, South Africa, 2010, 94.9%, G1 |
| VP7 | MG181727, *Homo sapiens*, Malawi, 2014 | 316 (1010-314) - 470 (437-476) | RDP 3.69E-17<br>GENECONV 1.84E-16<br>BootScan 8.96E-12<br>MaxChi 2.23E-08<br>Chimera 3.80E-08<br>SiScan 2.65E-12<br>3Seq 2.69E-23 | JQ069508, *Homo sapiens*, Canada, 2008, 98%, G1 | KJ753362, *Homo sapiens*, South Africa, 2003, 98.7%, G2 |
| VP7 | KX655511, *Homo sapiens*, Uganda, 2013 | 840 (823-851)- 1052 (1037-79) | RDP 1.20E-16<br>GENECONV 3.11E-15<br>BootScan 2.21E-16<br>MaxChi 6.12E-05<br>Chimera 6.98E-06<br>SiScan 2.52E-04<br>3Seq 3.36E-09 | EU839936, *Homo sapiens*, Bangladesh, 2005, 97.4%, G9 | KX655456, *Homo sapiens*, Uganda, 2013, 100%, G8 |
| VP7 | KJ753024, *Homo sapiens*, South Africa, 2009 | 1013 (985-59) - 130(121-143) | RDP 1.53E-16<br>GENECONV 2.32E-14<br>BootScan 1.27E-10<br>MaxChi 9.65E-04<br>Chimera 5.96E-04<br>SiScan 8.42E-13<br>3Seq 2.38E-16 | KM008672, *Homo sapiens*, India, 2010, 97.8%, G12 | D16344, *Homo sapiens*, 96.5%, Japan, 1977, G1 |
| VP7 | KJ751729, *Homo sapiens*, Ethiopia, 2010 | 857 (833-859) - 1019 (991-51) | RDP 4.0E-15<br>GENECONV 1.60E-10<br>BootScan 1.21EE-11 | KJ560447, *Homo sapiens*, USA, 2011, 99%, G3 | KP882703, *Homo sapiens*, Kenya, 2009, 99.3%, G1 |
|  | KP752817, *Homo sapiens*, Togo, 2010 |  | MaxChi 1.12E-03<br>Chimera 6.78E-04<br>SiScan 1.73E-03<br>3Seq 1.34E-05 |  |  |
| VP7 | MG181661, *Homo sapiens*, Malawi, 2013 | 590 (582-616) - 175 (170-181) | RDP 7.82E-30<br>GENECONV 7.33E-23<br>BootScan 5.06E-31<br>MaxChi 3.49E-21<br>Chimera 1.06E-25<br>3Seq 2.28E-47 | JQ069502, *Homo sapiens*, Canada, 2008, 96.6%, G1 | MG181859, *Homo sapiens*, Malawi, 2012, 99.5%, G2 |
| VP7 | KF170899, *Bos taurus*, India, 2010 | (71) - (448-506) | RDP 2.78 E-05<br>GENECONV 4.77 E-04<br>BootScan 4.86 E-04<br>MaxChi 1.02 E-03<br>Chimera 3.78 E-05<br>SiScan 7.21 E-09<br>3Seq 3.90 E-08 | EF199501, *Bos taurus*, India, 2001-2005, 97.3%, G6 | JX442784, *Bos taurus*, India, 2010, 99.5%, G6 |
| VP7 | KJ412857, *Homo sapiens*, Paraguay, 2009 | 772 (754-779) - 1014(988-51) | GENECONV 1.23E-19<br>BootScan 6.33E-20<br>MaxChi 2.56E-07<br>Chimera 4.38E-07<br>SiScan 2.01E-09<br>3Seq 3.36E-09 | KJ412858, *Homo sapiens*, Paraguay, 2009, 100%, G3 | KJ412802, *Homo sapiens*, Paraguay, 2009, 99.2%, G1 |
| VP7 | KC443034, *Homo sapiens*, USA, 2006<br>MG181727, *Homo sapiens*, Malawi, 2014 | 55 (994-60) - 291 (262-296) | RDP 1.66 E-30<br>GENECONV 5.07E-26<br>BootScan 2.32E-24<br>MaxChi 1.03E32<br>Chimera 1.10E-17<br>3Seq 3.36E-09 | GQ229044, *Homo sapiens*, India, 2005-2008, 97.4%, G2 | JX411970, *Homo sapiens*, India, 2011, 98.7%, G1 |
| VP7 | MG181771, *Homo sapiens*, Malawi, 2012 | 715 (707-722) - 950 (921-971) | RDP 2.65E-29<br>GENECONV 6.45E-26<br>BootScan 5.84E-17<br>MaxChi 1.70E-11<br>Chimera 2.36E-11<br>SiScan 4.78E-11<br>3Seq 3.55E-09 | KU356667, *Homo sapiens*, Bangladesh, 2013, 94.9%, G2 | KP222809, *Homo sapiens*, Mozambique, 2011, 99.6%, G1 |
| VP7 | KP752498, *Homo sapiens*, Togo, 2009 | 618 (605-641) - 1018 (1000-1020) | RDP 1.47E-28<br>GENECONV 4.49E-23<br>BootScan 2.78E-14<br>MaxChi 2.05E-14<br>Chimera 7.10E-19<br>SiScan 6.78E-39 | D86266, *Homo sapiens*, Japan, 1995, 95.7%, G3 | MG181661, *Homo sapiens*, Malawi, 2013, 98.8%, G1 |
| VP7 | AY261339, *Homo sapiens*, South Africa | 783 (774-791) - 14(1044-18) | RDP 3.82E-28<br>GENECONV 2.35E-25<br>BootScan 2.41E-27<br>MaxChi 5.44E-10<br>Chimera 2.33E-10<br>SiScan 3.15E-14<br>3Seq 3.36E-09 | GU567778, *Homo sapiens*, Russia, 2003-2009, 96.9%, G2 | JX458964, *Homo sapiens*, Argentina, 2001, 100%, G1 |
| VP7 | KX655489, *Homo sapiens*, Uganda, 2013 | 151 (132-155) - 316 (311-325) | RDP 2.14E-27<br>GENECONV 1.27E024<br>BootScan 3.15E-19<br>MaxChi 7.75E-09<br>Chimera 7.42E-09<br>SiScan 8.44E-20<br>3Seq 4.13E-03 | DQ146676, *Homo sapiens*, Bangladesh, 2003, 98.1%, G12 | JN232074, *Homo sapiens*, Brazil, 2004, 99.4%, G1 |
| VP7 | D86273, *Homo sapiens*, 1986 | 347 (339-353) - 726 (697-763) | RDP 1.70E-24<br>GENECONV 1.57E-22<br>BootScan 1.26E-17<br>MaxChi 7.23E-14<br>Chimera 1.37E-15<br>SiScan 1.73E-13<br>3Seq 6.15E-47 | AF450293, *Homo sapiens*, China, Unknown, 97.4%, G3 | D50121, *Homo sapiens*, 96.3%, Japan, 1992-1993, G2 |
| VP7 | KJ4122858, *Homo sapiens*, Paraguay, 2009 | 881 (859-914) - 1016 (991-54) | RDP 4.80E-14<br>GENECONV 3.04E-13<br>BootScan 3.81E-13<br>MaxChi 5.36E-03<br>Chimera 1.05E-02 | AB585923, *Homo sapiens*, Hong Kong, 2006, 98.5%, G3 | HQ425288, *Homo sapiens*, South Korea, 2004, 99.2%, G4 |
|  |  |  | SiScan 1.36E-04<br>3Seq 3.36E-09 |  |  |
| VP7 | KU714449, *Homo sapiens*, Malawi, 2000 | 825 (795-835) - 93 (85-93) | RDP 1.04E-13<br>GENECONV 4.64E-09<br>BootScan 1.71E-09<br>MaxChi 1.20E-10<br>Chimera 9.96E-11<br>SiScan 3.89E-10<br>3Seq 7.72E-08 | KM008635, *Homo sapiens*, India, 2009, 92.5%, G1 | KU714460, *Homo sapiens*, Malawi, 2000, 98.2%, G6 |
| VP7 | KJ412888, *Homo sapiens*, Paraguay, 2009 | 873 (823-890) - 1023 (1000-81) | RDP 2.17E-12<br>GENECONV 7.16E-11<br>BootScan 3.63E-12<br>MaxChi 5.48E-04<br>Chimera 4.30E-04<br>SiScan 8.10E-03<br>3Seq 1.12E-05 | D86275, *Homo sapiens*, China, 1986, 88%, G3 | JF490143, *Homo sapiens*, Australia, 2004, 99.3%, G1 |
| VP7 | KX655445, *Homo sapiens*, Uganda, 2013 | 1042 (1006-110) - 274 (269-287) | RDP 6.32E-12<br>GENECONV 5.98E-10<br>BootScan 8.41E-08<br>MaxChi 2.20E-06<br>Chimera 3.68E-08<br>SiScan 3.20E-04<br>3Seq 8.29E-11 | KJ560447, *Homo sapiens*, USA, 2011, 95.8%, G3 | KJ753326, *Homo sapiens*, Kenya, unknown, 94.9%, G8 |
| VP7 | D86277, *Homo sapiens*, China, 1989 | 335 (323-353)- 754 (710-762) | RDP 8.20E-20<br>GENECONV 2.21E-15<br>BootScan 1.81E-10<br>MaxChi 1.91E-12<br>Chimera 1.95E-13<br>SiScan 3.36E-13<br>3Seq 7.22E-27 | D86279, *Homo sapiens*, China, 1992, 89.6%, G3 | KP883055, *Homo sapiens*, Mali, 2009, 95.7%, G2 |
| VP7 | AF281044, *Homo sapiens*, Ireland, 1997-1999<br>GQ433992, *Homo sapiens*, Ireland, Unknown | 362 (346-366) - 589 (558-605) | RDP 1.32E-09<br>GENECONV 1.25E-02<br>BootScan 4.96E-05<br>MaxChi 2.75E-05<br>Chimera 1.50E-04<br>3Seq 3.36E-09 | AY866503, *Homo sapiens*, Thailand, 1995-1997, 98.9%, G9 | KM008633, *Homo sapiens*, India, 2009, 89%, G1 |
| VP7 | KU714449, *Homo sapiens*, Malawi, 2000 | 440 (421-446) - 604(641-704) | RDP 3.12E-10<br>GENECONV 7.36E-11<br>BootScan 1.56E-07<br>MaxChi 1.83E-03<br>Chimera 1.69E-03<br>3Seq 1.08E-12 | JQ069503, *Homo sapiens*, Malawi, 2000, 100%, G1 | KU714460, *Homo sapiens*, Malawi, 2000, 100%, G6 |
| VP7 | KX655444, *Homo sapiens*, Uganda, 2013 | 110 (88-114) - 191(183-197) | RDP 2.84E-08<br>GENECONV 3.00E-05<br>MaxChi3.11E-02<br>Chimera 3.11E-02<br>SiScan 1.01E-03<br>3Seq 8.25E-05 | KC579965, *Homo sapiens*, USA, 1978, 96.2%, G3 | AF254137, *Homo sapiens*, Ireland, 1997-1999, 93.9%, G1 |
| NSP1 | KP753174, *Sus scrofa*, South Africa, 2008,<br>KJ753184, *Sus scrofa*, South Africa, 2008,<br>KP752951, *Sus scrofa*, South Africa, 2008 | 1498 (1435-25) - 585(573-596) | RDP 7.45E-28<br>GENECONV 5.26E-27<br>BootScan 7.87E-25<br>MaxChi 8.49E-16<br>Chimera 5.73E-16<br>SiScan 9.95E-18<br>3Seq 1.09E-48 | KP752999, *Sus scrofa*, South Africa, 2009, 99.9%, A8 | KP753000, *Sus scrofa*, South Africa, 2009, 99.7%, A8 |
| NSP1 | KX655446, *Homo sapiens*, Uganda, 2013 | 485 (455-499)- 889 (868-914) | RDP 3.98E-25<br>GENECONV 2.55E-22<br>BootScan 2.53E-21<br>MaxChi 1.96E-14<br>Chimera 8.97E-15<br>SiScan 1.22E-19<br>3Seq 3.18E-09 | LC406825, *Homo sapiens*, Kenya, 2012, 95.9%, A2 | MG181277, *Homo sapiens*, Malawi, 2000, 96.5%, A1 |
| NSP1 | DQ199658, *Homo sapiens*, Australia, 2001 | 485 (440-517) - 942 (931-956) | RDP 2.25E-22<br>GENECONV 7.04E-20<br>BootScan 1.52E-21<br>MaxChi 2.09E-12<br>Chimera 1.10E-12<br>SiScan 6.14E-12<br>3Seq 2.61E-35 | KP883160, *Homo sapiens*, Malawi, 2008, 98.1%, A1 | DQ199657, *Homo sapiens*, Australia, 1997, 98.7%, A1 |
| NSP1 | KJ482257, *Sus scrofa*, Brazil, 2013 | 1534 (1490-44) - 888 (853-909) | RDP 3.24E-20<br>GENECONV 2.11E-19<br>BootScan 5.21E-10 | KJ482252, *Sus scrofa*, Brazil, 2013, 92.9%, A8 | KJ482255, *Sus scrofa*, Brazil, 2013, 99.9%, A8 |
|  |  |  | MaxChi 9.52E-18<br>SiScan 2.81E-21<br>3Seq 1.81E-29 |  |  |
| NSP1 | KX778616, *Homo sapiens*,<br>Dominican Republic, 2013 | 1147 (1127-1155) -<br>1462 (1433-5) | RDP 1.14E-19<br>GENECONV 4.51E-17<br>BootScan 8.47E-16<br>MaxChi 1.54E-11<br>Chimera 1.93E-12<br>SiScan 9.28E-13<br>3Seq 3.18E-09 | KJ752290, *Homo sapiens*, Zambia,<br>2009, 96.3%, A1 | KX363351, *Sus scrofa*,<br>Vietnam, 2012, 95.9%,<br>A8 |
| NSP1 | KJ482254, *Sus scrofa*,<br>Brazil, 2013 | 534 (499-559) -<br>886 (849-911) | RDP 7.72E-18<br>GENECONV 1.84E-16<br>BootScan 3.89E-14<br>MaxChi 2.53E-10<br>Chimera 2.27E-10<br>SiScan 1.55E-08<br>3Seq 8.01E-16 | KJ482247, *Sus scrofa*,<br>Brazil, 2013, 100%,<br>A8 | KJ482256, *Sus scrofa*,<br>Brazil, 2013, 100%, A8 |
| NSP1 | KX632348, *Homo sapiens*,<br>Uganda, 2013 | 580 (508-605) -<br>844 (829-874) | RDP 1.91E-10<br>GENECONV 3.85E-10<br>BootScan 1.02E-06<br>MaxChi 2.32E-04<br>Chimera 2.29E-04<br>SiScan 6.75E-08<br>3Seq 3.18E-09 | KP752785, *Homo sapiens*, Ethiopia,<br>2010, 94.9%, A1 | KP882434, *Homo sapiens*, Ghana, 2009,<br>92.8%, A2 |
| NSP1 | KP752999, *Sus scrofa*,<br>South Africa, 2009 | 1394 (1368-31) -<br>601 (552-631) | RDP 1.38E-06<br>BootScan 5.20E-05<br>MaxChi 2.04E-08<br>Chimera 2.74E-08<br>SiScan 3.04E-06<br>3Seq 2.78E-12 | KF835942, *Homo sapiens*, Hungary,<br>2005, 93.9%, A8 | Unknown, closest =<br>KP753000 *Sus scrofa*,<br>South Africa, 2009, A8 |
| NSP1 | JX040426, *Homo sapiens*,<br>India, 2009 | 662 (559-677) -<br>842 (783-884) | RDP 8.72E-03<br>GENECONV 1.35E-05<br>BootScan 4.19E-02<br>MaxChi 1.57E-04<br>Chimera 5.50E-05<br>SiScan 4.50E-03<br>3Seq 6.62E-07 | KU292525, *Homo sapiens*, India, 2014,<br>98.4%, A11 | KP882478, *Homo sapiens*, Ghana, 2008,<br>95%, A11 |
| NSP1 | KP753000, *Sus scrofa*,<br>South Africa, 2009 | 1389 (1359-41) -<br>602 (504-631) | RDP 4.27E-03<br>BootScan 4.30E-03<br>MaxChi 1.12E-04<br>Chimera 1.52E-05<br>SiScan 3.90E-04<br>3Seq 5.58E-05 | JQ309141, *Equus caballus*, United<br>Kingdom, 1975,<br>87.2%, A8 | KF835942, *Homo sapiens*, Hungary, 2005,<br>92.8%, A8 |
| NSP1 | KJ482254, *Sus scrofa*,<br>Brazil, 2013 | 1534 (1461-44) -<br>533 (499-909) | GENECONV: 1.50E-13<br>BootScan: 5.96E-10<br>MaxChi: 3.77E-08<br>Chimera: 6.43E-08<br>SiScan: 1.88E-11<br>3Seq: 1.46 E-24 | KJ482247, *Sus scrofa*,<br>Brazil, 2013, 99.1%,<br>A8 | KJ482255, *Sus scrofa*,<br>Brazil, 2013, 100%,<br>A8 |
| NSP2 | KJ753657, *Homo sapiens*,<br>South Africa, 2004<br>KM026663, *Homo sapiens*,<br>Brazil, 2009<br>KM026664, *Homo sapiens*,<br>Brazil, 2009 | (0) - (219 -278) | RDP 2.01E-03<br>GENECONV 1.6E-04<br>BootScan 4.07E-06<br>MaxChi 3.51E-03<br>Chimera 3.30E-03<br>SiScan 1.90E-07<br>3Seq 1.57E-04 | KM026633, *Homo sapiens*, Brazil, 2003,<br>98.6%, N1 | KJ918915, *Homo sapiens*,<br>Hungary, 2012, 99.5%,<br>N1 |
| NSP2 | MF18783, *Homo sapiens*,<br>USA, 2011 | 1015 (1002-57) -<br>286 (273-297) | GENECONV 1.60E-14<br>BootScan 1.55E-14<br>MaxChi 6.46E-04<br>Chimera 3.74E-06<br>SiScan 4.37E-08<br>3Seq 5.75E-22 | KJ752146, *Homo sapiens*, South Africa,<br>2011, 99.7%, N1 | KX655480, *Homo sapiens*, Uganda, 2012,<br>99.3%, N2 |
| NSP2 | MF18783, *Homo sapiens*,<br>USA, 2011 | 1015 (1002-57) -<br>286 (273-297) | GENECONV 1.60E-14<br>BootScan 1.55E-14<br>MaxChi 6.46E-04<br>Chimera 3.74E-06<br>SiScan 4.37E-08<br>3Seq 5.75E-22 | KJ752146, *Homo sapiens*, South Africa,<br>2011, 99.7%, N1 | KX655480, *Homo sapiens*, Uganda, 2012,<br>99.3%, N2 |
| NSP2 | AB779641, *Sus scrofa*,<br>Thailand, 2008 | 870 (833-921) -<br>216 (150-396) | RDP 6.30E-05<br>GENECONV 4.67 E-03<br>BootScan 1.24 E-03 | KX363374, *Sus scrofa*, Vietnam,<br>2012, 96.5%, N1 | JF796719, *Sus scrofa*,<br>South Korea, 2006, 100%,<br>N1 |
|  |  |  | MaxChi 1.28 E-03<br>Chimaera 6.20 E-03<br>3Seq 1.48 E-05 |  |  |
| NSP4 | MF58092, *Homo sapiens*,<br>China, 2013 | 599 (546-616) -<br>(745-94) | RDP 1.64 E-11<br>GENECONV 2.38 E-08<br>BootScan 1.14 E-09<br>MaxChi 1.01 E-09<br>Chimaera 4.58 E-10<br>SiScan 4.77 E-18<br>3Seq 6.43 E-21 | JF813107, *Homo sapiens*, China, 2008,<br>97.9%, E1 | MG781050, *Homo sapiens*, Thailand, 2011,<br>73.1%, E1 |
| NSP4 | AB361290, *Homo sapiens*,<br>India, 2006 | 624 (548-635) -<br>726 (712-50) | RDP 1.66 E-09<br>GENECONV 4.38 E-08<br>BootScan 6.96 E-07<br>MaxChi 1.96 E-03<br>Chimaera 2.49 E-05<br>3Seq 1.15 E-12 | AB326338, *Homo sapiens*, India, 1989,<br>95.8%, E2 | Unknown, closest =<br>FJ492834, *Sus scrofa*,<br>Ireland, E9 |
| NSP4 | AB326966, *Homo sapiens*,<br>India, 2001 | 605 (564-614) -<br>675 (670-46) | RDP 2.77 E-09<br>GENECONV 1.10 E-10<br>BootScan 2.26 E-02<br>MaxChi 2.54 E-05<br>Chimaera 1.63 E-03<br>3Seq 8.89 E-09 | FJ685615, *Homo sapiens*, India, 1992,<br>98.2%, E1 | Unknown, closest =<br>MG181929, *Homo sapiens*, Malawi, 2012, E2 |
| NSP5 | M33608, *Homo sapiens*,<br>1993 | 619 (505-624) -<br>668 (665-182) | RDP 1.44 E-05<br>GENECONV 2.64 E-06<br>BootScan 4.1 E-05<br>MaxChi 4.54 E-02<br>Chimaera 3.70 E-02<br>SiScan 2.49 E-05<br>3Seq 6.61 E-04 | MG996138, *Homo sapiens*, Singapore,<br>2016, 95%, H2 | JQ358774, *Homo sapiens*,<br>India, 2011, 100%, H3 |

**Supplementary Table 2.** Amino acid changes observed in segment 9 (VP7) recombination events observed in more than one isolate. Changes occurring in predicted epitope regions are noted.

| Gene recombinant | Accession Number | Position | Major | Minor | Epitope region? |
| --- | --- | --- | --- | --- | --- |
| G6-G6 | KF170899 | 40 | M | T | - |
|  |  | 46 | I | T | - |
|  |  | 48 | V | T | - |
|  |  | 116 | V | I | - |
| G3-G1 | KJ751729 | 281 | V | I | Yes |
|  |  | 303 | V | I | Yes |
|  |  | 309 | A | V | Yes |
|  |  | 318 | N | D |  |
| G9-G1 | AF281044 | 135 | S | C | - |
|  |  | 162 | D | E | - |
|  |  | 163 | R | W | - |
|  |  | 169 | V | D |  |
|  |  | 170 | N | I |  |
|  |  | 176 | L | Q |  |
|  |  | 180 | G | E | Yes |
| G2-G1 | KC443034 | 9 | I | T | - |
|  |  | 10 | F | I | - |
|  |  | 21 | S | Y | - |
|  |  | 25 | T | S | - |
|  |  | 26 | I | V | - |
|  |  | 28 | N | R | - |
|  |  | 29 | P | I | - |
|  |  | 31 | A | D | - |
|  |  | 32 | P | Y | - |
|  |  | 35 | F | Y | - |
|  |  | 41 | I | F | - |
|  |  | 42 | A | V | - |
|  |  | 43 | L | A | - |
|  |  | 44 | M | L | - |
|  |  | 45 | S | F | - |
|  |  | 47 | F | L | - |
|  |  | 48 | G | T | - |
|  |  | 49 | R | K | - |
|  |  | 50 | T | A | - |
|  |  | 56 | Y | N | - |
|  |  | 65 | A | T | - |
|  |  | 68 | T | S | - |
|  |  | 72 | S | R | - |
|  |  | 73 | G | E | - |
|  |  | 75 | S | V | - |

**Supplementary Figure 1.**
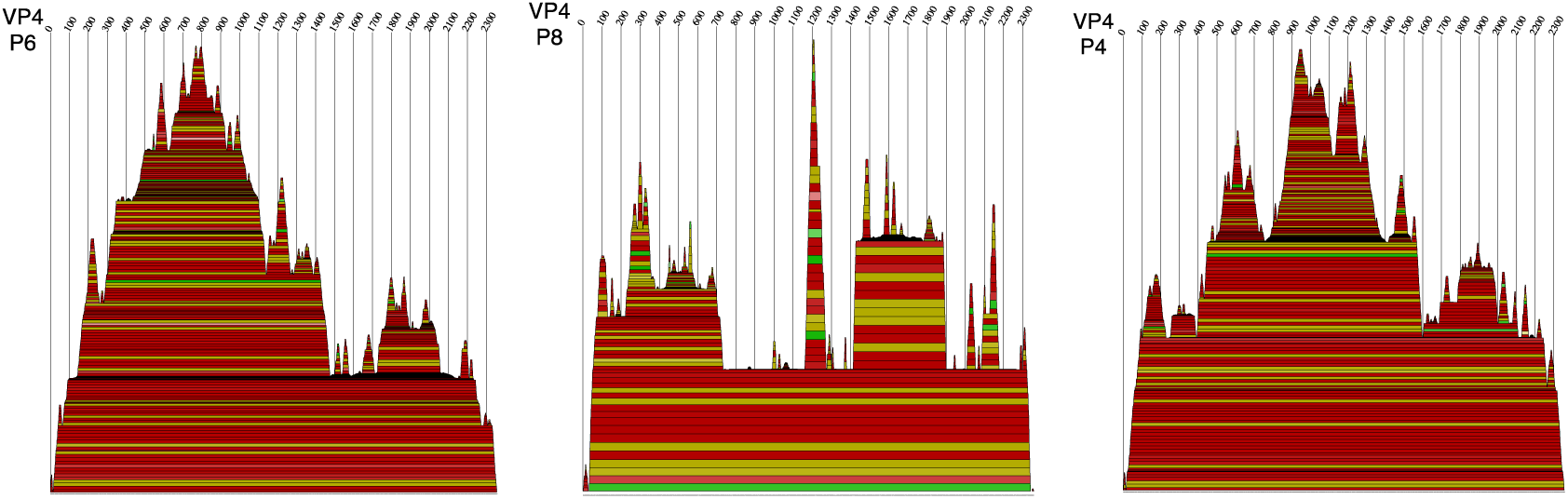
P6, P8, and P4 consensus RNA secondary structure mountain plots. Mountain plots showing consensus secondary structure of VP4 genotypes P6, P8, and P4. Lighter colors indicate lower probabilities of base-pairing. Peaks correspond to hairpin-loops, plateus to loops, and slopes to helices. Plot was generated using RNAalifold in the ViennaRNA package.

**Supplementary Figure 2.**
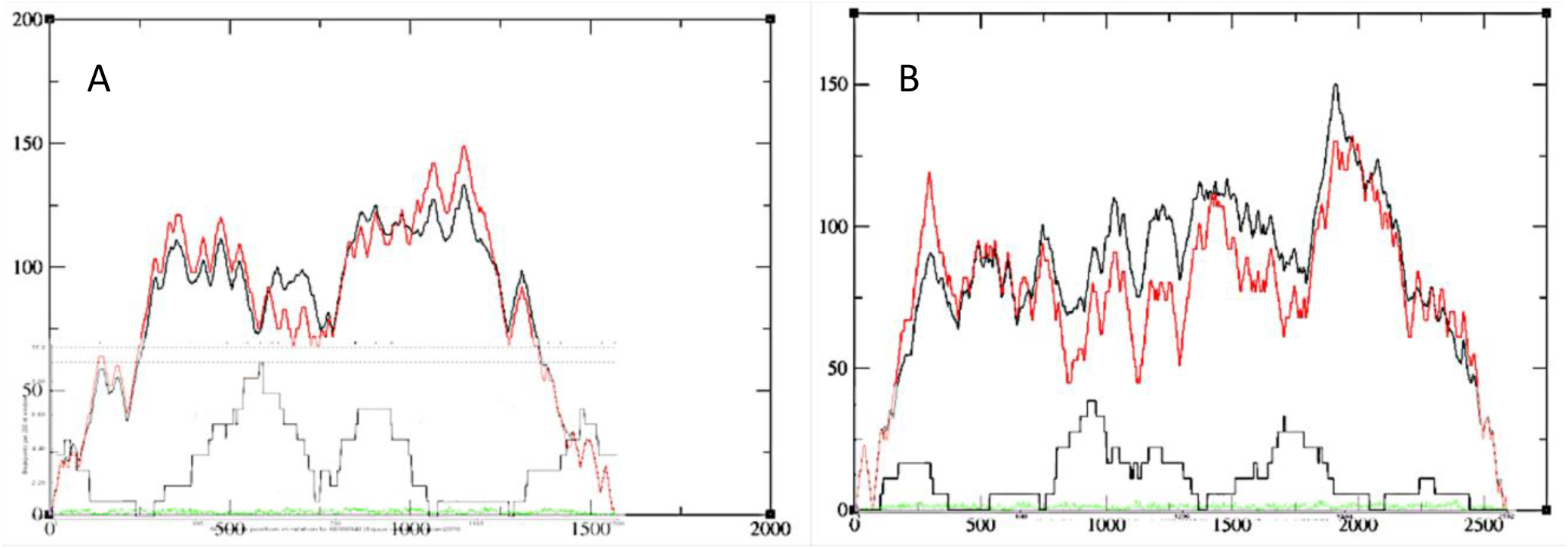
Consensus mountain plots of A) NSP1 and B) VP3 with recombination breakpoint distribution plot in gray.

